# KDM5 inhibition offers a novel therapeutic strategy for the treatment of *KMT2D* mutant lymphomas

**DOI:** 10.1101/2020.06.30.177477

**Authors:** James A Heward, Lola Konali, Annalisa D’Avola, Karina Close, Alison Yeomans, Martin Philpott, James Dunford, Tahrima Rahim, Ahad F Al Seraihi, Jun Wang, Koorosh Korfi, Shamzah Araf, Sameena Iqbal, Findlay Bewicke-Copley, Emil Kumar, Darko Barisic, Maria Calaminici, Andrew Clear, John Gribben, Peter Johnson, Richard Neve, Jessica Okosun, Udo Oppermann, Ari Melnick, Graham Packham, Jude Fitzgibbon

## Abstract

Loss-of-function mutations in *KMT2D* are a striking feature of the germinal centre (GC) lymphomas, resulting in decreased H3K4 methylation and altered gene expression. We hypothesised that inhibition of the KDM5 family, which demethylates H3K4me3/me2, would re-establish H3K4 methylation and restore the expression of genes repressed upon loss of *KMT2D*. KDM5-inhibition increased H3K4me3 levels and caused an anti-proliferative response *in vitro*, which was markedly greater in both endogenous and CRISPR-edited *KMT2D* mutant DLBCL cell lines, whilst tumour growth was inhibited in *KMT2D* mutant xenografts *in vivo*. KDM5-inhibition reactivated both KMT2D-dependent and -independent genes, resulting in diminished B-cell receptor signalling and altered expression of BCL2 family members, including BCL2 itself, allowing it to synergise with agents targeting these pathways. KDM5-inhibition may offer an effective therapeutic strategy for ameliorating *KMT2D* loss-of-function mutations in GC-lymphomas.

**Statement of significance:** We detail a novel way of reverting the effects of loss-of-function mutations in the histone methyltransferase *KMT2D* by inhibiting the KDM5 demethylase family, increasing levels of H3K4me3 and restoring expression of KMT2D regulated genes.

## Introduction

Although epigenetic dysregulation is a feature of most cancers, few are as strikingly dependent as GC-lymphomas. The vast majority of Follicular Lymphoma (FL) tumours harbour loss-of-function mutations in *KMT2D* (80%) alongside mutations in *CREBBP* (60%) and *EZH2* (25%) (1-4), with *KMT2D* also frequently mutated (30%) within the GC B-cell (GCB) subtype of Diffuse Large B-cell Lymphoma (DLBCL) (5-7). The majority of *KMT2D* mutations in GC-lymphomas are truncating, arise early during tumour development, and are often bi-allelic (1,8-10), yet despite their frequency, no therapies targeting these mutations have been reported.

The histone methyltransferase KMT2D (ENSG00000167548; formerly MLL2 or Mll4) is a member of the KMT2 family of methyltransferases (KMT2A-H) which catalyse the mono-, di- and tri-methylation of Histone 3 Lysine 4 (H3K4) (11). These modifications are generally associated with active transcription, with H3K4me1 predominantly located at enhancers and H3K4me3 at active and poised promoters (12). KMT2D has preferential mono-methyltransferase activity and deposits H3K4me1 at enhancers, although it also acts as the central structural-component of the COMPASS-like multi-protein complex and is required for the correct recruitment of other enzymes including the histone acetyltransferases EP300/CREBBP and the H3K27me3 demethylase KDM6A (UTX) (13,14).

Loss of *Kmt2d* has been demonstrated to decrease H3K4me1/me2 deposition, alter gene expression and to co-operate with *Bcl2* overexpression in VavP-*Bcl2* mice, increasing proliferation within the GC and driving lymphomagenesis. Germline *KMT2D* mutations are also the predominant cause of Kabuki syndrome, a developmental disorder with defects in B-cell development but no apparent increase in GC-lymphoma prevalence (15,16), highlighting that *KMT2D* mutations are likely to co-operate with other lesions to cause GC-lymphomas.

H3K4 methylation levels are also regulated by the Lysine Specific Demethylase (KDM) families LSD1 and KDM5, which demethylate H3K4me1 to H3K4me0 and H3K4me3/me2 to H3K4me1 respectively. The KDM5 family utilises α-ketoglutarate as a substrate and contains four members; *KDM5A (JARID1A/RBP2), KDM5B (JARID1B/PLU1), KDM5C (JARID1C/SMCX)* and *KDM5D (JARID1D/SMCY)*. The KDM5 family has essential roles in regulating gene expression in a variety of contexts, and although mutations of KDM5 genes are rare, KDM5A and KDM5B have been implicated as potential therapeutic targets due to their upregulation in several cancers (17) and apparent role as drivers of metastasis and drug resistance (18-20).

In this report, we hypothesised that KDM5-inhibition would re-establish H3K4 methylation and restore the expression of genes deregulated upon loss of *KMT2D*. Using several different KDM5-inhibitors (KDM5i), we demonstrate that KDM5-inhibition has strong anti-proliferative and cytotoxic activity on GCB-DLBCL cell lines, likely through a combination of regulating BCR-signalling and the expression of BCL2 family members. Critically, KDM5-inhibition sensitivity appears to be dependent on the presence of *KMT2D* mutations, suggesting that KDM5-inhibition may offer a targeted therapy for *KMT2D* mutant GC-lymphomas.

## Results

### KDM5-inhibition increases global H3K4me3 levels in GC-lymphoma cells

To assess whether the KDM5 family was a suitable therapeutic target for GC-lymphomas, we first quantified the expression of the four KDM5 isoforms (*KDM5A*-*D*) in DLBCL cell lines, primary FL (ICGC (21)) and DLBCL (ICGC/TCGA) biopsies and normal GC B-cells (BLUEPRINT (22)) (Figure 1a+b). *KDM5A* and *KDM5C* were highly expressed in all the samples whilst expression of the Y-linked *KDM5D* was restricted to male derived cell lines (Figure 1a). Protein expression was confirmed for KDM5A, KDM5C and KDM5D by western blot analysis (Supplementary Figure 1a).

**Figure 1.**
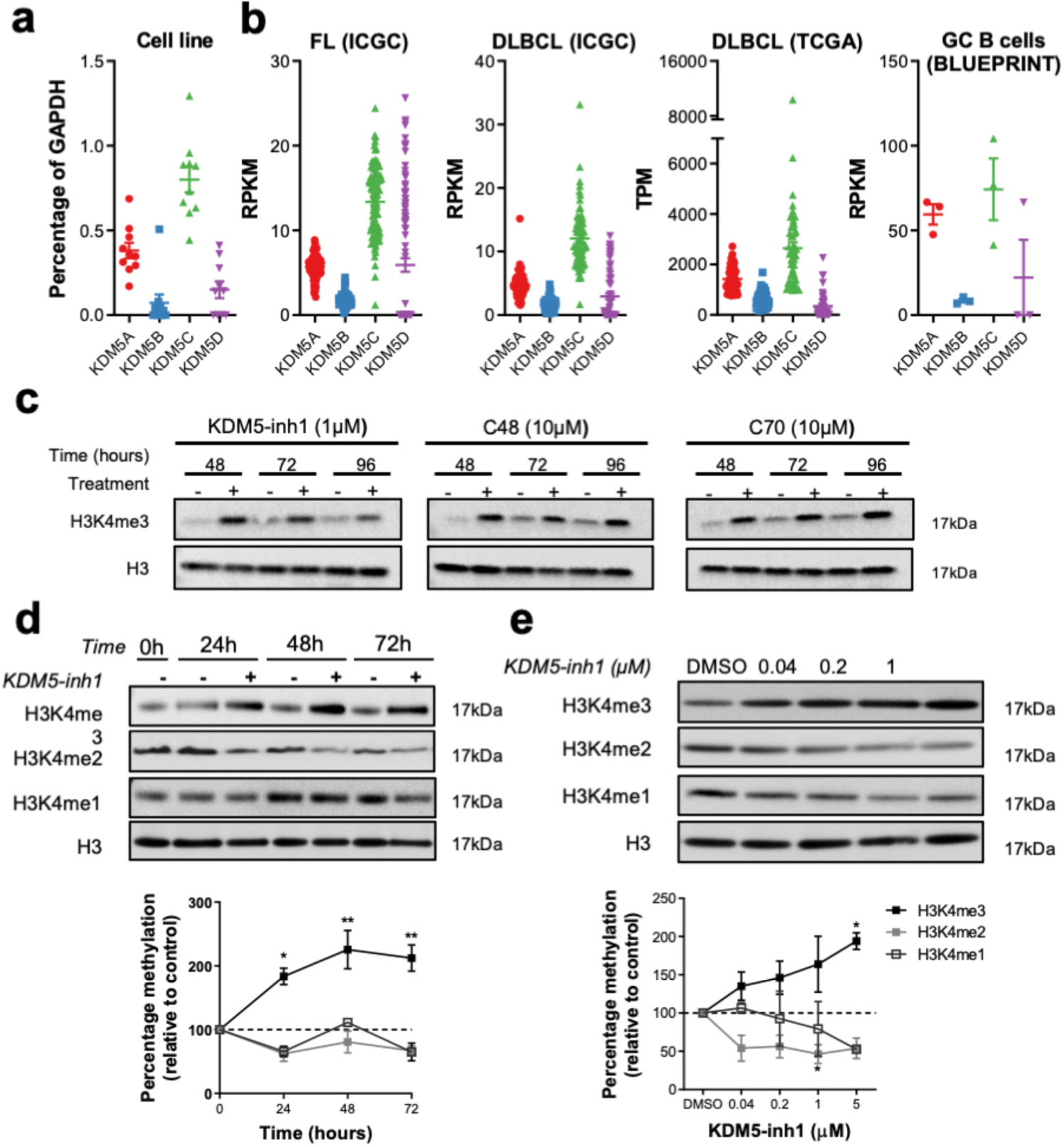
KDM5-inhibition increases H3K4me3 levels in DLBCL cell lines. **(a)** The expression of the four *KDM5* family members (*KDM5A-D*) was examined by qRT-PCR in 10 DLBCL cell lines and normalised to the expression of GAPDH. Data are the mean ± SEM of three independent experiments. **(b)** KDM5 family member expression was examined by RNA-seq in publicly available datasets of FL (ICGC, n=97 (21)) and DLBCL (TCGA n=48, ICGC n=74) patients, plus healthy GC B-cells (BLUEPRINT (22)). RPKM = Reads Per Kilobase Million, TPM = Transcripts Per Million. **(c)** SU-DHL-6 cells were treated with 1µM KDM5-inh1 or 10µM Compound-48 and KDM5-C70 for 48h, 72h and 96h, followed by western blot analysis of H3K4me3 levels relative to H3. The SU-DHL-6 cell line was **(d)** treated with DMSO or 1μM KDM5-inh1 for increasing lengths of time and **(e)** for 48h with DMSO or increasing concentrations of KDM5-inh1. The upper panels display representative western blots for H3K4me3/me2/me1 and H3. The lower panel displays the quantification of western blots relative to H3. Data are the mean ± SEM of 3 independent experiments. Statistical significance was determined using an ANOVA with a Dunnett’s post-test versus untreated control, where * P<0.05 and ** P<0.01.

We then examined the effect of three individual KDM5i on H3K4 methylation; KDM5-inh1 (Patent no. WO 2014/131777 A1 – EpiTherapeutics/Gilead (23)), Compound-48 (Constellation Pharmaceuticals (24)) and KDM5-C70 (25). All three KDM5i increased H3K4me3, with KDM5-inh1 the most potent and Compound-48 and KDM5-C70 requiring concentrations around 10-fold higher to induce similar increases in H3K4me3 (Figure 1c). In GC-lymphoma cell lines, KDM5-inh1 induced time- and concentration-dependent increases in H3K4me3, alongside modest decreases in H3K4me1/me2 (Figure 1d+e; Supplementary Figure 1b+c), without altering KDM5A and KDM5C protein levels (Supplementary Figure 1d+e). KDM5-inh1 had no effect on histone marks mediated by the two closest related KDM families, KDM4 (H3K9me3/H3K36me3) and KDM6 (H3K27me3) (17), indicating that KDM5-inh1 is specific for KDM5 (Supplementary Figure 2a+b). KDM5-inhibition also increased H3K4me3 in primary FL cell-suspensions (n=8), with H3K4me3 increased to a greater degree in both *KMT2D* mutant cell lines and cell-suspensions, versus WT, at 48h (Supplementary Figure 2c+d).

**Figure 2.**
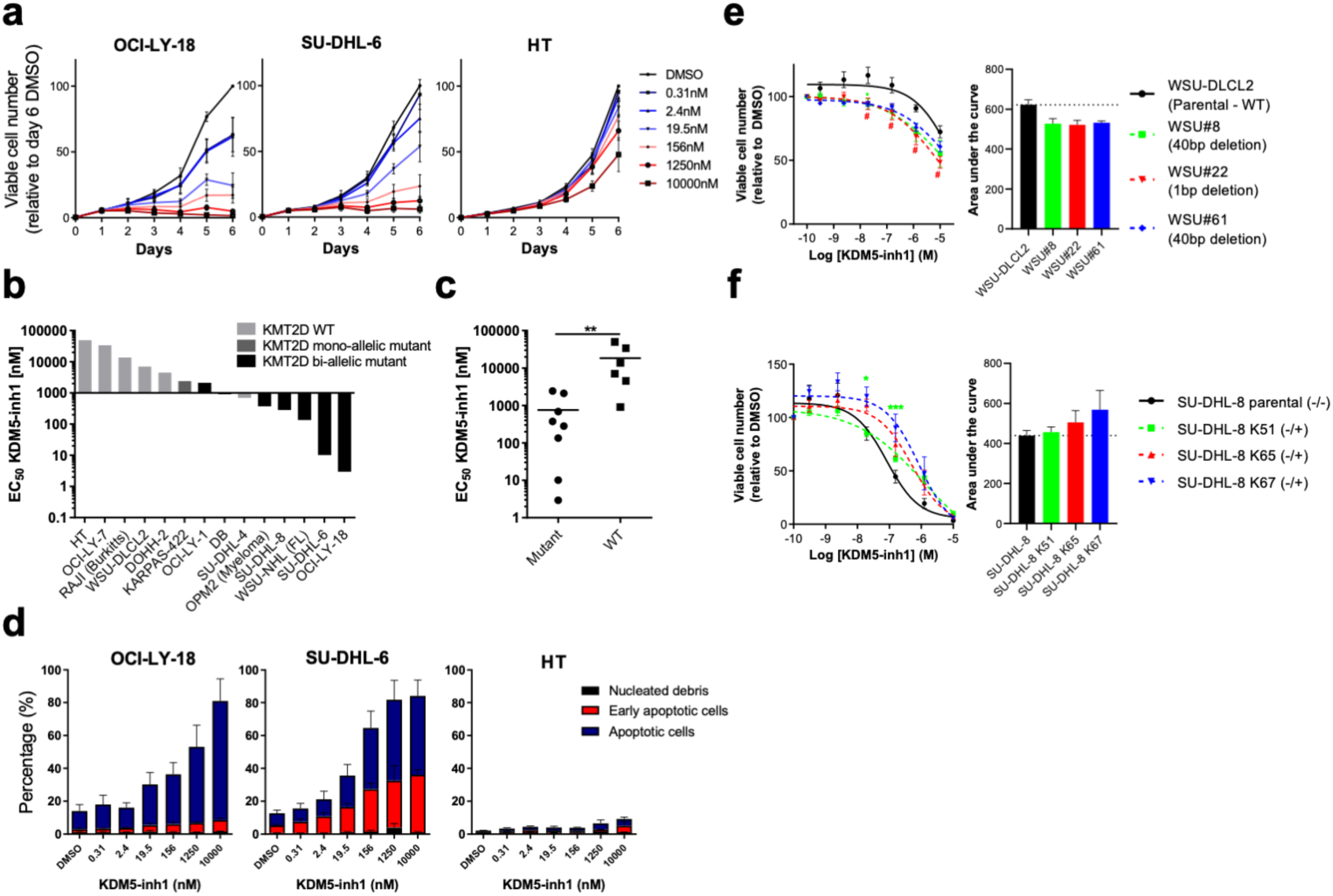
KDM5-inhibition reduces the proliferation of *KMT2D* mutant cell lines. DLBCL, FL, myeloma and Burkitt’s lymphoma cell lines were treated with DMSO or increasing concentrations of KDM5-inh1, and viable cells quantified **(a)** every day up to 6 days for OCI-LY-18, SU-DHL-6 and HT cells **(b)** and after 5 days for all cell lines, with EC_50_ values for *KMT2D* WT and mutant cell lines displayed in a waterfall plot. **(c)** Dot plot showing the significantly lower EC_50_ values for *KMT2D* mutant cell lines. Statistical significance was determined by Mann Whitney U test, where ** P<0.01. **(d)** Induction of apoptosis was quantified in OCI-LY-18, SU-DHL-6 and HT cells treated with DMSO or increasing concentrations KDM5-inh1 for 5 days. Viable cell counts from **(e)** WSU-DLCL2 cells and 3 *KMT2D* mutant clones or **(f)** parental SU-DHL-8 cells and 3 corrected clones treated with DMSO or increasing concentrations of KDM5-inh1 for 5 days. Data are the mean ± SEM of 3-7 independent experiments. Statistical significance was calculated using a two-way ANOVA with a Dunnett’s post-test, where */# P <0.005 and *** P < 0.001.

### KDM5-inhibition has selective cytostatic and cytotoxic activity on *KMT2D* mutant cell lines

We next examined the cytostatic effect of KDM5-inhibition on an extended panel of cell lines. KDM5-inh1 had a varied impact upon proliferation after five days, with some cell lines insensitive and others displaying strikingly low EC_50_ values (e.g. OCI-LY-18 = 3nM, SU-DHL-6 = 10nM; Figure 2a+b; Supplementary Figure 3a). Compound-48 and KDM5-C70 were less potent, although reduced proliferation was observed in SU-DHL-6 and OCI-LY-18, the cell lines most sensitive to KDM5-inh1 (Supplementary Figure 3a). Grouping of the cell lines by *KMT2D* mutation status revealed that KDM5-inh1 had a significantly greater anti-proliferative effect upon *KMT2D* mutant cell lines (Mann-Whitney U, P value = 0.003; Figure 2b+c), with eight out of nine of the most sensitive harbouring *KMT2D* mutations. The majority of cell lines examined (6/8) displayed lower EC_50_ values after 10 days of treatment than at five days (Supplementary Figure 3b+c), indicating that KDM5-inhibition has sustained anti-proliferative activity in lymphoma cells. Furthermore, quantification of DNA content and Annexin/7-AAD staining indicated that the most sensitive cell lines were undergoing apoptosis following KDM5-inhibition (Figure 2d; Supplementary Figure 3d-f).

**Figure 3.**
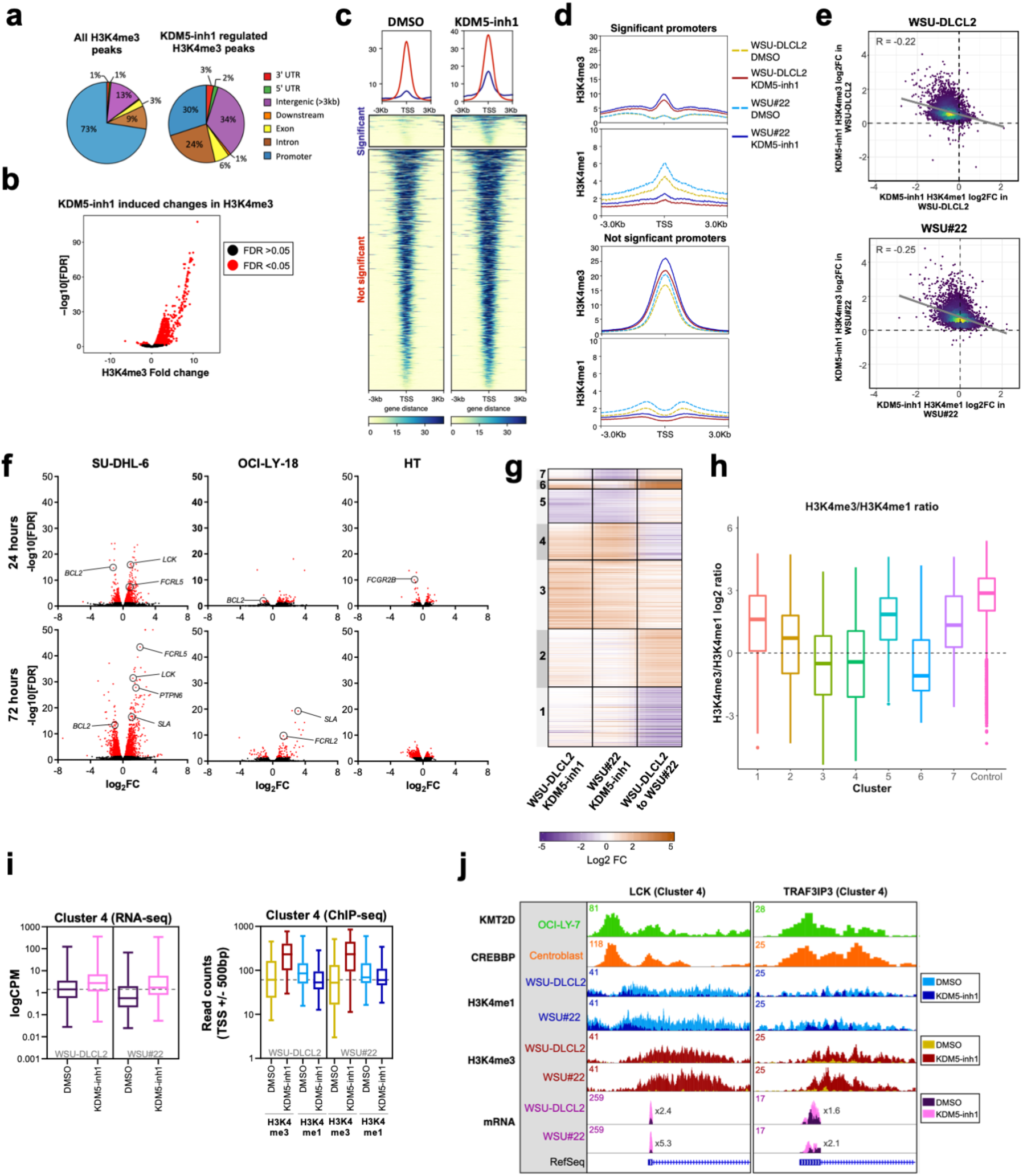
Epigenetic and transcriptomic characterisation of KDM5-inhibition. **a)** Genomic locations of H3K4me3 peaks identified by ChIP-seq in cells treated with DMSO (left) or 1μM KDM5-inh1 (right) for 72h. **(b)** KDM5-inhibition induced changes in H3K4me3, with significantly changed peaks displayed in red. **(c)** Heatmaps of ChIP-seq data showing difference in H3K4me3 levels between promoters significantly altered (blue) or otherwise (red) in SU-DHL-6 cells treated with DMSO or 1µM KDM5-inh1 for 72h. **(d)** Spatial plots showing distribution of H3K4me1 and H3K4me3 at promoters with significantly altered H3K4me3 by KDM5-inh1 or otherwise, in WSU-DLCL2 (yellow/red) and WSU#22^-/+^ (light/dark blue) cells treated with DMSO (yellow/light blue) or KDM5-inh1 (red/dark blue). **(e)** Plots showing broad increases in H3K4me3 and reductions in H3K4me1, quantified by ChIP-seq, at the TSS (+/- 500bp) of H3K4me3+ genes in WSU-DLCL2 and WSU#22^-/+^ cells treated with 1μM KDM5-inh1. The Pearson’s correlation co-efficient is indicated on each plot. **(f)** Volcano plots indicating DE genes in SU-DHL-6, OCI-LY-18 and HT cells treated with 1μM KDM5-inh1 for 24h and 72h, with significant genes highlighted in red. **(g)** Heatmap showing log2FC values for 897 genes that were DE by either KDM5-inh1 or *KMT2D* loss, and clustered using K-means clustering. **(h)** H3K4me3 and H3K4me1 reads were counted for the promoters in each cluster, then divided (H3K4me3/H3K4me1) and log2 normalised to create a summary ratio for each. Control promoters were identified as being H3K4me3+ in WSU-DLCL2 cells but showing no alteration in mRNA expression or H3K4me3/H3K4me1 deposition in any of our analyses. **(i)** Boxplots showing RNA-seq logCPM values (left) and TSS read counts across ChIP-seq (right) datasets, of genes from Cluster Four. **(j)** ChIP-seq and RNA-seq tracks, centred on *LCK* and *TRAF3IP3*, from WSU-DLCL2 and WSU#22^-/+^ cells treated with KDM5-inh1 for 72h, plus ChIP-seq tracks of KMT2D (OCI-LY-7) (9) and CREBBP (centroblasts) (29) binding.

### Inducing and correcting *KMT2D* mutations by CRISPR alters KDM5-inhibition sensitivity

Since *KMT2D* mutant cells were more sensitive to KDM5-inhibition than WT cells (Figure 2b+c), we next tested whether inducing or correcting *KMT2D* mutations in cell lines would alter KDM5-inhibition sensitivity. Using CRISPR we introduced *KMT2D* mutations in two WT cell lines; WSU-DLCL2, the least sensitive t(14;18) positive cell line and HT, the least sensitive cell line overall. Three WSU-DLCL2 clones (#8, #22, #61; Supplementary Table 1) harbouring mono-allelic truncating mutations displayed reduced proliferation following KDM5-inhibition (Figure 2e), with an average decrease of 16% in area under the curve (AUC) values. Global levels of H3K4me3/me2/me1 appeared unaltered by *KMT2D* loss in untreated cells whilst KDM5-inhibition induced similar increases in H3K4me3 in mutant and WT cells (Supplementary Figure 4a+b). In contrast to WSU-DLCL2, the CRISPR-edited *KMT2D* mutant HT cells appeared intrinsically resistant to KDM5-inhibition, with no consistent changes in proliferation observed (Supplementary Figure 4c+d).

**Figure 4.**
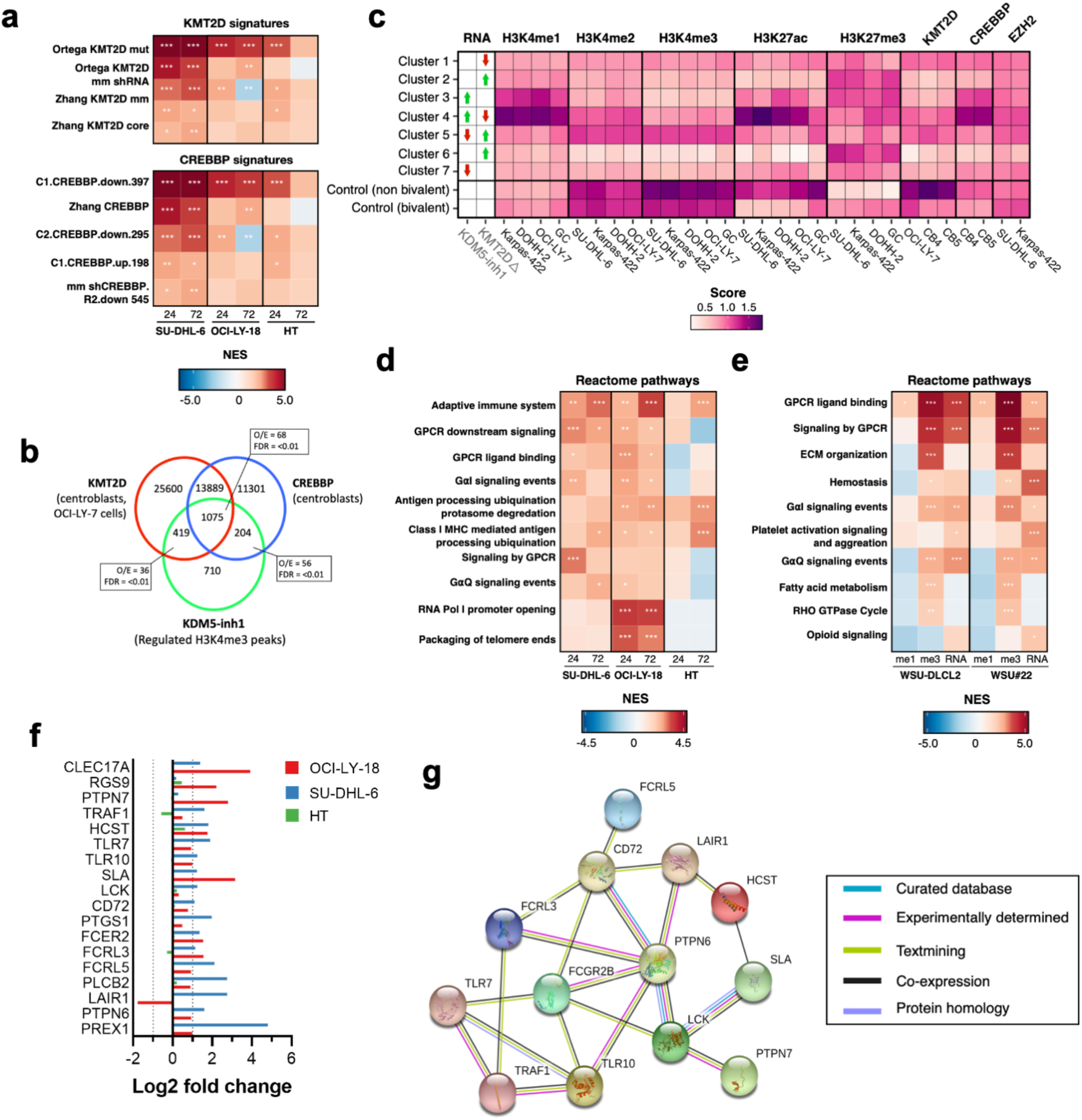
KDM5-inhibition regulates KMT2D target genes and BCR-signalling regulators. **(a)** Heatmap indicating normalized enrichments scores (NES) of KMT2D and CREBBP signatures in KDM5-inh1 treated cells, following GSEA of RNA-seq profiles using a manually curated database of B-cell signatures. **(b)** Overlap between KDM5-inhibition regulated regions in SU-DHL-6 and CREBBP (29) or KMT2D bound regions (9,10), with observed/expected and FDR values for the overlaps indicated. **(c)** Deeptools (58) was used to calculate summary scores at the promoters (TSS+/-500bp) of genes in each cluster (Figure 3f), plus non-bivalent (H3K4me3+/H3K27me3-) and bivalent (H3K4me3/H3K27me3+) control promoters, for ChIP-seq datasets of histone mark deposition (ENCODE/BLUEPRINT) and KMT2D (9,10), CREBBP (29) and EZH2/SUZH12 (34) binding. The overall direction of change in RNA expression, following KDM5i or *KMT2D* loss, is indicated for each cluster in the first two columns. **(d+e)** Heatmaps indicating NES following GSEA of the Reactome database in **(d)** RNA-seq profiles from KDM5-inh1 treated cells and **(e)** RNA-seq plus promoter H3K4me1 and H3K4me3 profiles from KDM5-inh1 treated WSU-DLCL2/WSU#22^-^ /+ cells. **(f)** Log2FC values of BCR-signalling regulators in SU-DHL-6, OCI-LY-18 and HT cells treated with KDM5-inh1. **(g)** String analysis (https://string-db.org/) showing the interaction network of identified BCR-signalling regulators.

CRISPR was also employed to correct the homozygous 1bp insertion (P648Tfs*2) that disrupts *KMT2D* in the KDM5-inhibition sensitive SU-DHL-8 cells, generating three clones where a single allele had been reverted to WT, two of which displayed increased global H3K4me1 (K51 and K65; Supplementary Figure 4e). All of these clones were more resistant to KDM5-inhibition with an average increase in AUC of 19% (Figure 2f), confirming that KDM5-inhibition sensitivity is altered by *KMT2D* mutations.

### KDM5-inhibition induces widespread increases in H3K4me3

We hypothesised that increased H3K4me3 levels would drive a gene expression programme responsible for the cytostatic and cytotoxic activity of KDM5-inhibition. H3K4me3 ChIP-seq identified 11158 H3K4me3 peaks in untreated SU-DHL-6 cells (Supplementary Table 2; Supplementary Figure 5a), with the majority (72.6%) located at gene promoters (Figure 3a). KDM5-inhibition increased the average peak size (Supplementary Figure 5b) and altered H3K4me3 levels at 2408 peaks, with 98% demonstrating increased H3K4me3 (Supplementary Table 2; Figure 3b). Only a third of these peaks overlapped with promoters (Figure 3a), suggesting that KDM5-inhibition may alter H3K4me3 deposition at both enhancers and promoters. This was confirmed by overlaying intergenic regions regulated by KDM5-inhibition with ChIP-seq data from GC-lymphoma cell lines (ENCODE) and primary GC B-cells (BLUEPRINT (22)), which showed that 84-95% overlapped with the enhancer-associated H3K4me1 mark (Supplementary Figure 5c+d). These intergenic regions also largely showed deposition of H3K4me3 and H3K27ac, indicating that the majority are active enhancers.

**Figure 5.**
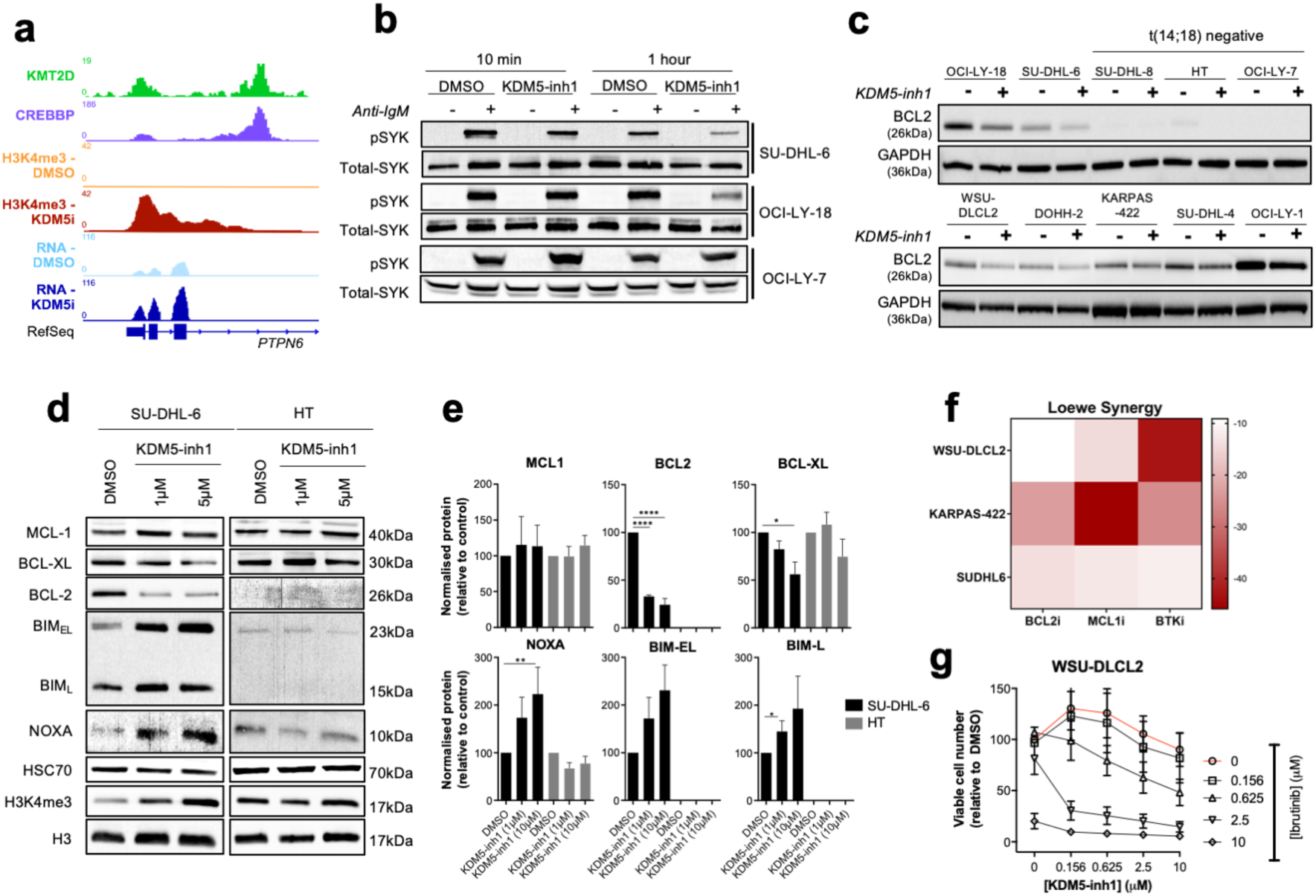
KDM5-inhibition alters the expression of BCR-signalling and apoptotic regulatory genes. **(a)** ChIP-seq and RNA-seq tracks, centred on the *PTPN6* promoter, from SU-DHL-6 cells treated with DMSO or 1μM KDM5-inh1 for 72h, plus ChIP-seq tracks of KMT2D (9) and CREBBP (29) binding in GC centroblasts. **(b)** Activation of the BCR-associated kinase SYK was investigated by western blot analysis in SU-DHL-6, OCI-LY-18 and OCI-LY-7 cells treated with DMSO or KDM5-inh1 for 72h, followed by anti-IgM F(ab’)_2_ antibody stimulation for 10 min or 1h. **(c)** Expression of BCL2 protein was examined by western blot in 10 DLBCL cell lines exposed to DMSO or 1μM KDM5-inh1 48h. **(d)** SU-DHL-6 and HT cells were treated with DMSO or 1μM and 5μM KDM5-inh1 for 5 days, with the cells re-seeded in fresh drug/media after 48h. The expression of BCL2 family members was investigated by western blot, with HSC70 used as a loading control. Western blots are representative of 3 independent experiments, with the quantification relative to HSC70 displayed in **(e)**. Statistical significance was determined using an ANOVA with a Dunnett’s post-test versus untreated control, where * P<0.05, ** P<0.01 and **** P < 0.0001. **(f)** SU-DHL-6, KARPAS-422 and WSU-DLCL2 cells were treated with increasing concentrations of KDM5-inh1 for 5 days, alongside increasing concentrations of S63845 (MCL1i), Venetoclax (BCL2i) for 2 days or Ibrutinib (BTKi) for 3 days. Viable cells were quantified and an overall Loewe synergy score calculated for each combination. **(g)** Plot showing viable cell numbers following KDM5-inh1 and Ibrutinib combinations in WSU-DLCL2 cells. Data are the mean ± SEM of 3 independent experiments.

We also noted that promoters significantly altered by KDM5-inhibition displayed basal levels of H3K4me3 that were significantly lower than the average promoter in SU-DHL-6 (Figure 3c). These promoters also displayed low levels of H3K4me3 in other cell lines (e.g. OCI-LY-7) and instead had higher levels of H3K4me1 (Supplementary Figure 5e+f). A low H3K4me3/H3K4me1 ratio has previously been described to mark promoters that are poised to respond to cellular signalling (26,27), and it is probable that the KDM5 family maintains a poised configuration at these promoters by preventing high levels of H3K4me3 deposition.

### KDM5-inhibition converts H3K4me1 to H3K4me3 at promoters

To understand how *KMT2D* mutations alter H3K4me1 and H3K4me3, and the influence this has on the response to KDM5-inhibition, we focused on WSU-DLCL2 clone #22 (WSU#22^-/+^), where we had engineered a heterozygous 1bp deletion (P95Qfs*35) that is typical of the *KMT2D* mutations seen in GC-lymphomas (Figure 2e). We first examined global changes in H3K4me3/me1 by ChIP-seq, and observed moderate changes in H3K4me1 (1333 altered peaks; 62.3% decreased) between untreated WSU-DLCL2 and WSU#22^-/+^ cells, with H3K4me3 minimally affected (49 altered peaks) (Supplementary Figure 6a-c; Supplementary Table 2). The response to KDM5-inhibition was more dramatic, with H3K4me3 deposition broadly increased (>99% of sites) in WSU-DLCL2 and WSU#22^-/+^ cells (4604/3244 peaks) while there was a predominant reduction (>80%) in H3K4me1 levels (2469/3130 peaks) (Supplementary Figure 6a-c; Supplementary Table 2).

**Figure 6.**
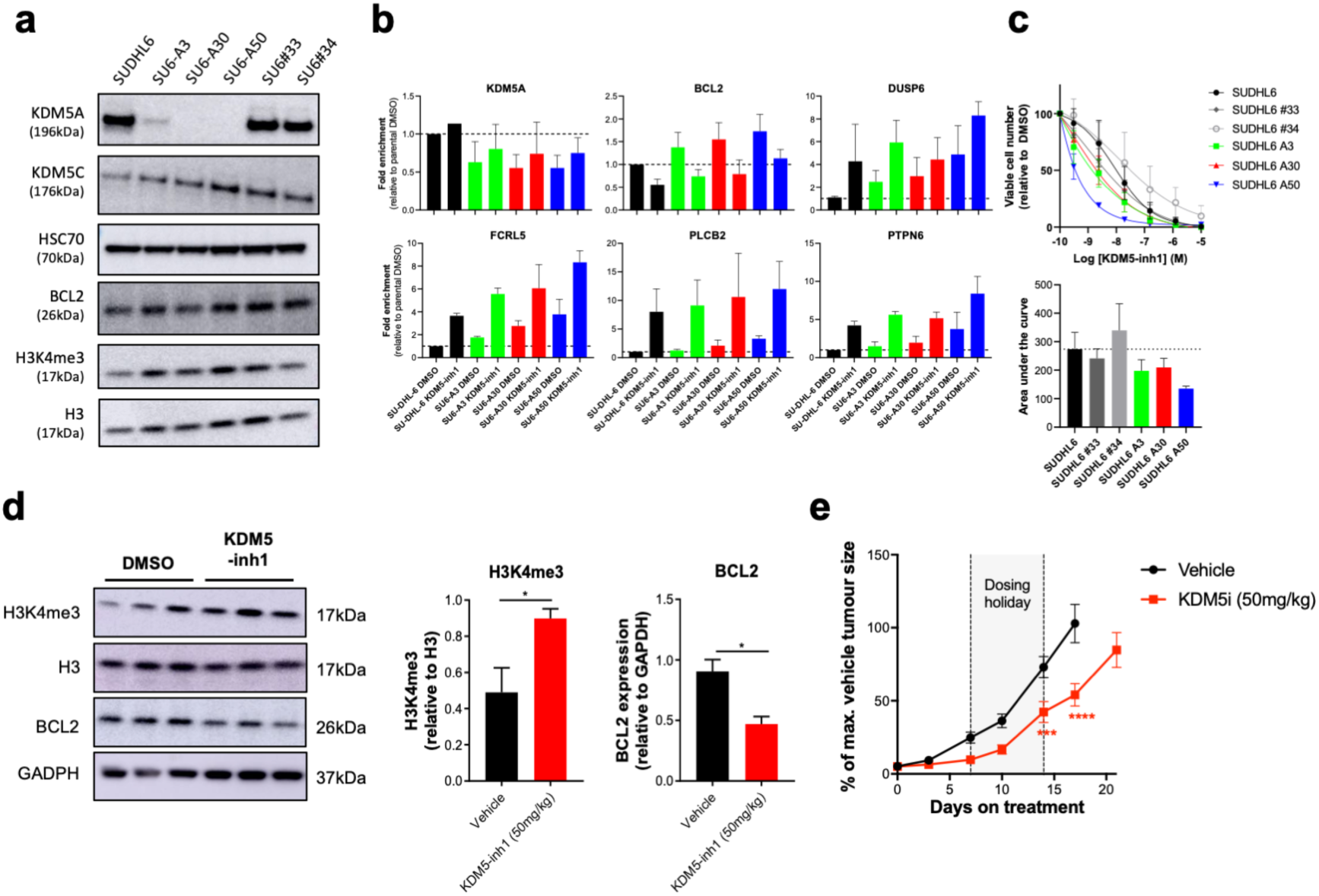
KDM5-inhibition likely acts through multiple isoforms and is efficacious *in vivo*. **(a)** Western blot showing loss of KDM5A in three homozygous knockout clones (SU6-A3, A30, A50) compared to parental and WT controls (SU6#33, #34), alongside expression of KDM5C, BCL2 and H3K4me3 levels. **(b)** qRT-PCR analysis of *KDM5A* and KDM5-inhibition target genes in SU-DHL-6 and KDM5A knockout clones exposed to DMSO or 1μM KDM5-inh1 for 72h. Data are the mean ± SEM of two independent experiments. **(c)** SU-DHL-6 and KDM5A knockout clones were exposed to DMSO or increasing concentrations of KDM5-inh1 for five days, and viable cell numbers quantified. Concentration response curves are shown in the upper panel and AUC values in the lower panel. Data are the mean ± SEM of three independent experiments. **(d)** Global levels of H3K4me3 in tumours from mice treated with vehicle or 50mg/kg KDM5-inh1 for 1 week (n=3) were quantified by western blot and normalised to H3, whilst BCL2 levels were quantified and normalised to GAPDH. Quantification of the western blots is displayed on the right-hand side. **(e)** Activity of 50mg/kg KDM5-inh1 on the growth of SU-DHL-6 xenografts, in comparison to vehicle treated mice. Data are the mean ± SEM of 10 individual mice, except in the vehicle group where one mouse was removed due to insufficient tumour growth (<300mm^3^). Statistical significance was calculated using a two-way ANOVA with a Dunnett’s post-test, where *** P <0.001 and **** P < 0.0001.

We identified 10,259 promoters that were marked by H3K4me3 but not significantly altered by KDM5-inhibition in WSU-DLCL2 or WSU#22^-/+^ cells, and 1958 promoters with significantly altered H3K4me3 following KDM5-inhibition. As before (Supplementary Figure 5e+f), we found the majority of promoters to display a typical high H3K4me3/H3K4me1 ratio whilst the significantly altered promoters showed an inverse low H3K4me3/H3K4me1 ratio (Figure 3d; Supplementary Figure 6d). Across all promoters KDM5-inhibition reduced H3K4me1 and increased H3K4me3 (Figure 3d+e), although the degree of change was more striking in the H3K4me1 high/H3K4me3 low group (Figure 3d) and suggests that KDM5-inhibition activates promoters by converting H3K4me1 into H3K4me3.

### KDM5-inhibition induces moderate changes in gene expression

Genes differentially expressed (DE; FDR <0.05, log2FC > 1 or <-1) by KDM5-inhibition were identified by RNA-seq analysis of two sensitive (SU-DHL-6 and OCI-LY-18) and one insensitive cell line (HT) treated with 1μM KDM5-inh1 for 24h or 72h. Overall, a greater number of DE genes were observed at 72h versus 24h in all the cell lines tested (Figure 3f; Supplementary Table 3) with the impact on expression most striking in the SU-DHL-6 cell line (147 and 545 DE genes). Modest changes in expression occurred in the other sensitive cell line OCI-LY-18 (52 and 83 DE genes) and the insensitive cell line HT (13 and 95 DE genes). In all conditions, with the exception of HT 72h, the majority of DE genes were upregulated, whilst there was a greater overlap between the two sensitive cell lines (Supplementary Figure 7a-c).

Focusing on SU-DHL-6 and comparing our KDM5-inhibition RNA- and ChIP-seq data at 72h, we observed that promoter H3K4me3 correlated with gene expression (r=0.28) to a greater extent than enhancer H3K4me3 levels (r=0.04; versus nearest gene) (Supplementary Figure 7d). This was more pronounced when selectively examining upregulated genes (0.44 vs 0.02) and indicates that KDM5-inhibition activates gene expression through promoters rather than enhancers, whilst gene downregulation may be an indirect consequence downstream of H3K4me3 deposition. Overall, these results indicate that KDM5-inhibition has a relatively modest impact upon gene expression despite inducing widespread increases in H3K4me3.

### KDM5-inhibition regulates KMT2D dependent and independent genes

We next compared RNA-seq profiles between WSU-DLCL2 and WSU#22^-/+^ cells and identified 445 DE genes, while parallel KDM5-inhibition led to 309 and 339 changes in gene expression, which included 141 common transcripts (Supplementary Figure 7e+f). In total, 897 genes were either DE between WSU-DLCL2 and WSU#22^-/+^ or following KDM5-inhibition, which were divided into seven discrete groups using K-mean clustering (Figure 3g; Supplementary Table 3).

The majority of genes (71%) were regulated by either KMT2D (e.g. Clusters One and Two) or KDM5-inhibition alone (e.g. Cluster Three). We also identified two clusters (Clusters Four and Five) where changes in gene expression accompanying the P95Qfs*35 mutation were effectively reversed following KDM5-inhibition. Cluster Four contained genes which were downregulated by *KMT2D* loss (mean log2FC = - 0.87) but upregulated by KDM5-inhibition in both WT and mutant cells (mean log2FC = 0.92 vs 1.75), including the cell-cycle regulator *CDKN1A* and several BCR-signalling regulators (*LCK, TRAF3IP3, PRKCB, FCGR2B*). An inverse-relationship was observed in Cluster Five, where genes were upregulated by *KMT2D* loss (mean log2FC = 0.31) but downregulated by KDM5-inhibition in WSU-DLCL2 and WSU#22^-/+^ cells (mean log2FC = -1.0 vs -1.1) (Figure 3g), including the apoptotic-regulator *BCL2*.

We next analysed how levels of promoter H3K4 methylation may regulate gene expression in these clusters. Clusters upregulated by KDM5-inhibition (Three, Four and Six) exhibited low basal H3K4me3/high H3K4me1, whereas the remaining clusters had a typical high H3K4me3/H3K4me1 ratio (Figure 3h, Supplementary Figure 7g). Although KDM5-inhibition reduced H3K4me1 and increased H3K4me3 within all the clusters, its effect was most notable upon Clusters Three and Four, where it altered the H3K4me3/H3K4me1 ratio to the extent that levels of H3K4me3 surpassed H3K4me1 (Figure 3i+j, Supplementary Figure 7g). Cluster Five in contrast displayed minimal changes in H3K4me3, supporting our previous observation that KDM5-inhibition may indirectly downregulate gene expression. Across all clusters however, *KMT2D* loss induced minimal changes to promoter H3K4me1/me3 (Supplementary Figure 7g), indicating that *KMT2D* mutations may not regulate gene expression through H3K4 methylation.

### KDM5-inhibition regulates KMT2D and CREBBP target genes

To further test whether KDM5-inhibition regulates KMT2D target genes, we used Gene Set Enrichment Analysis (GSEA) (28) to compare our two RNA-seq series with a manually-curated database of lymphoma and B-cell signatures, including signatures derived from patient cohorts, *in vitro* analyses and conditional mouse models of KMT2D and CREBBP loss (9,10,29-31). All four datasets generated in two recent lymphoma *KMT2D* studies (9,10) were significantly enriched in both series, as were 28 signatures associated with CREBBP (Figure 4a; Supplementary Figure 8a; Supplementary Table 3+4) and the HDAC3i BRD3308 (Supplementary Figure 8b), recently proposed as a targeted therapy to reverse the effects of CREBBP loss (30,31). Moreover, these CREBBP/KMT2D signatures were also enriched in our SU-DHL-6 and WSU-DLCL2/WSU#22^-/+^ H3K4me3 ChIP-seq data (Supplementary Figure 8a+c; Supplementary Table 4+5), whilst the 2408 regions regulated by KDM5-inhibition in SU-DHL-6 significantly overlapped with binding of KMT2D and CREBBP, with 62% of the regions bound by KMT2D, 53% by CREBBP and 45% by both (Figure 4b; Supplementary Figure 8d). In contrast, we detected modest enrichment for signatures associated with *EZH2* mutations and EZH2i (32,33), and limited overlap between KDM5-inhibition regulated regions and EZH2/SUZ12 binding (34)(Supplementary Figure 8a-d).

We next investigated the enrichment of a range of histone marks and epigenetic-regulators in our previously defined gene clusters in WSU-DLC2/WSU#22^-/+^ (Figure 3g), including H3K27ac, H3K27me3, KMT2D and CREBBP, in GC-lymphoma cell lines (ENCODE) and GC B-cells (BLUEPRINT (22)). In agreement with our earlier observations linking CREBBP to KDM5 regulated genes (Figure 4a+b), Cluster Four displayed levels of H3K27ac and CREBBP binding that were noticeably higher than cany other cluster, including Cluster Three which is regulated by KDM5-inhibition but not KMT2D loss (Figure 4c). Levels of KMT2D binding conversely did not appear to be predictive of KDM5-inhibition response (Figure 4c; Supplementary 8e). Since recent publications indicate that the major consequences of KMT2D loss may occur through altering EP300 and KDM6A recruitment (13,14), we propose that *KMT2D* mutations sensitise cells to KDM5i by altering the recruitment of other epigenetic enzymes (e.g. EP300/CREBBP), thereby repressing the expression of a subset of genes with an atypical epigenetic profile and a high dependency for H3K27ac, which can be reactivated through KDM5-inhibition converting H3K4me1 into H3K4me3.

### KDM5-inbition upregulates regulators of BCR-signalling

Pathway analysis of the genes associated with altered H3K4me3 in SU-DHL-6 revealed pathways highly relevant to lymphoma biology, with the most enriched terms including “Hematologic cancer”, “Adaptive immune system” and several pathways related to BCR-signalling, whilst pathways related to GPCR signalling were predominantly enriched in our H3K4me3 analysis of KDM5-inh1 treated WSU-DLCL2/WSU#22 cells (Supplementary Figure 8a+f; Supplementary Table 5). Similarly, GSEA (28) of our RNA-seq data identified the pathway “Adaptive immune system” as being strongly enriched across all conditions, whilst pathways related to GPCR-signalling were strongly enriched in SU-DHL-6, OCI-LY-18 and WSU-DLCL2/WSU#22^-/+^ cells (Figure 4d+e; Supplementary Table 4), indicating that KDM5-inhibition may regulate B-cell signalling. This was further supported by our observation that KDM5-inhibition primarily targets genes with high levels of promoter H3K4me1 (Supplementary Figure 5e-f), which has been described as a signature of signal-responsive genes (26,27), whilst BCR-signalling regulators were identified within the KDM5-inhibition and KMT2D regulated genes in Cluster Four (Figure 3g-j).

Amongst the negative-regulators of BCR-signalling induced by KDM5-inh1 was the tyrosine phosphatase SHP-1 (*PTPN6*; Figure 4f) (35,36), which is regulated by KMT2D (9) and CREBBP (29), and subject to low-frequency mutations and silencing in lymphoma (37), alongside a range of receptors able to recruit and activate SHP-1 including *FCGR2B, FCRL3/5, CD72* and *LAIR1* (Figure 4g). KDM5-inhibition increased H3K4me3 and reduced H3K4me1 levels across the *PTPN6* promoter in SU-DHL-6 cells, without altering H3K27ac, and upregulated SHP-1 expression (Figure 5a; Supplementary Figure 9a+b). Increased promoter H3K4me3 levels were observed for *PTPN6* and other BCR-signalling regulators in primary FL cell-suspensions following KDM5-inhibition (Supplementary Figure 9c), while these BCR-signalling regulators were also induced by Compound-48 and KDM5-C70 in SU-DHL-6 cells (Supplementary Figure 9d), indicating that these genes are specifically regulated by the demethylase activity of the KDM5 family.

### KDM5-inhibition results in a more rapid curtailment of BCR-signalling

To determine whether the increased expression of negative-regulators such as SHP-1 (35,36) altered BCR-signalling, the phosphorylation of SYK, a proximal kinase activated following BCR engagement, was examined in the IgM^+^ SU-DHL-6 and OCI-LY-18 cells (Supplementary Figure 10a) pre-treated with DMSO or KDM5-inh1 for 72h and then stimulated with anti-IgM F(ab’)_2_. KDM5-inh1 pre-treatment reduced levels of SYK phosphorylation at later time points (1-4h) following sIgM engagement, whilst the initial induction of SYK phosphorylation at 10 minutes was unaffected by KDM5-inhibition (Figure 5b, Supplementary Figure 10b+c). Effects on sIgM were selective since KDM5-inhibition did not affect surface expression of sIgM (Supplementary Figure 10a) or intracellular calcium release (Supplementary Figure 10d+e). By contrast, we observed KDM5-inhibition to have no impact upon SYK phosphorylation in the KDM5-inhibition insensitive anti-IgM responsive OCI-LY-7 cells (Figure 5b; Supplementary Figure 10b+c) and the KDM5-inhibition insensitive HT cells, which had extremely high constitutive levels of BCR-signalling and were unresponsive to anti-IgM stimulation (data not shown) (38). The kinetics of signalling without alterations of sIgM expression suggested a more rapid curtailment in BCR-signalling in KDM5-inhibition sensitive cells, consistent with the expected consequence of increasing the expression of regulators such as SHP-1 (39).

### KDM5-inhibition modulates the expression of BCL2 family members

Amongst the downregulated genes, we observed reduced expression of the anti-apoptotic BCL2. All three KDM5i tested consistently reduced BCL2 protein expression in t(14;18) positive cell lines (Figure 5c; Supplementary Figure 11a), although sensitivity to KDM5-inhibition and the BCL2i Venetoclax varied (r=0.38, p=0.31), indicating that response to KDM5-inhibition is not solely dependent on BCL2 (Supplementary Figure 11b+c). The mechanism of BCL2 downregulation appeared to be an indirect effect of KDM5-inhibition, as we observed no clear changes in H3K4me3/me1 or H3K27ac across the *BCL2* promoter (Supplementary Figure 9a) or at enhancers contained within *BCL2* or the IGH locus (data not shown), consistent with earlier observations that downregulated genes do not correlate with H3K4me3 deposition (Supplementary Figure 7d+g).

We next analysed the expression of BCL2 alongside other family members in SU-DHL-6 and HT cells treated with KDM5-inh1 for two and five days (OCI-LY-18 cells were not examined due to high levels of drug-induced cell death). Minimal changes were observed in the insensitive HT cells, however decreased BCL2 and BCL-XL expression, alongside increasing expression of the pro-apoptotic NOXA, BIM_L_ and BIM_EL_, were observed at day five in SU-DHL-6 (Figure 5d+e). These changes preceded the onset of apoptosis at day two (Supplementary Figure 11d+e), whilst KDM5-inhibition also reduced *BCL2* and *BCL-XL* mRNA expression in primary FL cell-suspensions (Supplementary Figure 9c). Overall, these data indicate that KDM5-inhibition shifts the balance of BCL2 family members towards a pro-apoptotic response in sensitive cells.

### KDM5-inhibition synergises with MCL1 and BTK inhibitors

Given the ability of KDM5-inhibition to regulate BCR-signalling and BCL2 family members, we next tested whether KDM5-inhibition could synergise with the BH3 mimetics Venetoclax (BCL2i) (40) and S63845 (MCL1i) (41) and the BTKi Ibrutinib (42), which are all under clinical-investigation for GC-lymphomas. Although we were unable to detect synergy in SU-DHL-6 cells due to their high response to KDM5-inhibition alone, we observed KDM5-inh1 to highly synergise with S63845 in KARPAS-422 (*KMT2D*^-/+^) and with Ibrutinib in WSU-DLCL2 (*KMT2D*^+/+^) cells (Figure 5f+g; Supplementary Figure 12, Supplementary Table 7). The synergy with the MCL1i S63845 is likely explained by KDM5-inhibition downregulating the expression of the two other major negative regulators of apoptosis, BCL2 and BCL-XL, whilst the synergy with ibrutinib could be due to altered expression of BCR-signalling regulators and/or BCL2 family members.

### Loss of *KDM5A* alone does not alter proliferation or survival

We next used CRISPR to examine how KDM5A and KDM5C regulate previously identified KDM5-inhibition target genes and determine whether any KDM5 isoform alone is essential for lymphoma survival. *KDM5A* and *KDM5C* knockout clones were successfully isolated from *KMT2D* WT WSU-DLCL2 cells (Supplementary Figure 13a) however we were only able to generate *KDM5A* knockout clones from *KMT2D* mutant SU-DHL-6 cells (Figure 6a). Loss of *KDM5A/KDM5C* in WSU-DLCL2 or *KDM5A* in SU-DHL-6 cells had minimal impact on H3K4me3 levels (Figure 6a; Supplementary Figure 13a) or upon cell proliferation or survival (data not shown). *KDM5A* knockout did upregulate the expression of several KDM5 target genes in SU-DHL-6 (e.g. *FCRL5, DUSP6*), although to a lesser extent than KDM5-inhibition (Figure 6b).

We next analysed whether the anti-proliferative response to KDM5-inhibition was altered in *KDM5A/KDM5C* knockout cells, reasoning that it would be blunted if KDM5-inhibition primarily functioned through either isoform alone, and found instead that silencing of *KDM5A* and *KDM5C* both increased sensitivity to KDM5-inhibition in SU-DHL-6 and WSU-DLCL2 (Figure 6c; Supplementary Figure 13b). Whilst it is possible that losing or inhibiting KDM5C alone may be lethal in *KMT2D* mutant cells, we believe it is more likely that multiple isoforms must be inhibited to robustly induce gene expression and an anti-proliferative response, which is supported by previous reports of redundancy in the KDM5 (43,44) and other KDM families (45,46), global H3K4me3 levels remaining stable in our single isoform knockout models and *KDM5A* knockout only partially activating KDM5-inhibition regulated genes.

### KDM5-inh1 has *in vivo* activity against *KMT2D* mutant xenografts

In order to test the *in vivo* efficacy of KDM5-inh1, *KMT2D* mutant SU-DHL-6 cells were xenografted subcutaneously into NOD/SCID mice. Mice were orally administered vehicle, 50mg/kg KDM5-inh1 daily or 10mg/kg Ibrutinib (positive control) for 21 days, with a dosing-holiday scheduled between days 8-14 for the KDM5-inh1 group after preliminary experiments indicated that this regime would be efficacious and tolerable. KDM5-inh1 was well tolerated throughout the study, with weight loss <20% and no other signs of toxicity observed (Supplementary Figure 14a-c). Levels of H3K4me3 were variable at the study endpoint, although increased H3K4me3 and reduced BCL2 expression were observed in the tumours of mice treated with KDM5-inh1 for seven days (Figure 6d; Supplementary Figure 14d). After seven days of treatment, tumour growth inhibition (TGI) of 65% was observed for the KDM5-inh1 group, and while the tumours partially recovered during the dosing-holiday, TGI values of 54-66% were maintained until day 17 when the vehicle group was sacrificed (Figure 6e).

## Discussion

The *KMT2* methyltransferases are one of the most highly disrupted gene families across cancer (11), most notably within the GC-lymphomas where 80% of FL (1-4) and 30% of GCB-DLBCL cases harbour *KMT2D* mutations (5-7), alongside mutations in other epigenetic-regulatory genes including *CREBBP* (29,30,47-49) and *EZH2* (50-52). The potential of precisely targeting these mutations has recently been established by the development of EZH2i, with these compounds partially selective towards *EZH2* mutant FL patients in phase II clinical trials (33,53,54). The observation that *CREBBP* mutant lymphomas are HDAC3-dependent has also presented another potential therapeutic target (30,31), and hints that pharmacological inhibition of antagonistic enzymes may be an effective strategy for targeting loss-of-function epigenetic mutations. Despite the high frequency of *KMT2D* mutations in GC-lymphomas and other malignancies, no means of therapeutically targeting these lesions has been reported. We therefore examined whether inhibiting the KDM5 family could ameliorate the loss of *KMT2D* by stabilising H3K4 methylation and restoring the expression of genes normally regulated by *KMT2D*.

KDM5-inhibition increased global levels of the promoter-associated H3K4me3 and induced significant cytostatic and cytotoxic responses in *KMT2D* mutant cell lines *in vitro* and tumour growth inhibition *in vivo*. In this report we describe two mechanisms through which KDM5i function. Firstly, we observed increased expression of negative-regulators of B-cell signalling, including *PTPN6*, resulting in the more rapid curtailment of BCR-signalling in cells treated with KDM5i. Secondly, KDM5-inhibition induced striking reductions in BCL2 expression in t(14;18) positive cell lines alongside altering the expression of other BCL2 family members. Although pharmacologically targeting BCL2 alone appears to have modest single-agent activity in GC-lymphomas (55,56), the simultaneous modulation of pro- and anti-apoptotic regulators, alongside the loss of survival signals from the BCR, seems likely to drive the induction of apoptosis triggered by KDM5-inhibition. Encouragingly, our data indicate that KDM5i are able to synergise with therapeutics agents targeting these pathways, although further *in vivo* studies are required to establish the therapeutic window of KDM5-inhibition, both alone and in combination, and whether it impacts upon normal B-cell functions. Our data is also consistent with prior reports of significant redundancy within the KDM5 (43,44), and other KDM families (45,46), and indicates that inhibition of multiple KDM5 members may be required to achieve a therapeutic response, which should be considered in the further development of KDM5i.

KDM5-inhibition sensitivity was confirmed to be highly dependent on *KMT2D* by generating and correcting *KMT2D* mutations in WT and mutant cell lines respectively, with alteration of a single allele capable of shifting the response to KMD5-inhibition. Our epigenetic and transcriptomic analyses revealed that KDM5-inhibition activates both KMT2D-dependent and -independent gene networks, and in particular targets promoters marked by high levels of H3K4me1. High H3K4me1 levels may act to maintain these promoters in a poised configuration, and have previously been described to mark the promoters of stimuli-responsive genes with roles in signal transduction (26,27). This is consistent with our observation of KDM5-inhibition increasing the expression of BCR-signalling regulators, while *KMT2D* mutations have been shown to alter B-cell signalling by preventing the upregulation of negative-regulators following CD40-stimulation (10).

Despite the evidence presented here that KDM5i can reactivate KMT2D target genes, by increasing H3K4me3 at the expense of H3K4me1, KDM5-inhibition does not directly reverse the epigenetic consequences of losing the mono-methyltransferase activity of KMT2D. Indeed inhibition of LSD1, the direct antagonist of the methyltransferase activity of KMT2D, has previously been shown to be ineffective in lymphoma, suggesting that restoring H3K4me1 alone is not sufficient to restore KMT2D-reglated genes (57). One potential explanation worth considering is the ability of KMT2D to recruit the H3K27 acetyltransferases EP300/CREBBP and demethylase KDM6A (13,14), which would imply that loss of H3K4me1 is only one part of a wider epigenetic disruption induced by *KMT2D* mutations.

Our data indicate that KMT2D-dependant genes upregulated by KDM5-inhibition (Cluster Four) display strikingly high levels of H3K27ac and CREBBP-binding in addition to a low H3K4me3/H3K4me1 ratio. Further studies are required to establish whether mutations in *KMT2D* alter the recruitment of EP300/CREBBP to these promoters, and while the extent to which epigenetic mutations overlap in lymphoma is a key outstanding question, our data indicates that this subset of genes is likely to be disrupted by multiple mutations. Systematically identifying genes regulated by multiple epigenetic-mutations may be one way to distil their key targets and determine the most effective therapeutic-agents and combinations to target these lesions.

In summary, this report establishes the potential of KDM5-inhibition as a targeted therapy for GC-lymphomas that is able to reactivate the expression of genes normally regulated by KMT2D. In particular, the increased expression of negative-regulators of B-cell signalling results in a curtailment of pro-survival signals and decreases the expression of BCL2 and other BCL2 family proteins. Notably, the response to KDM5-inhibition appears to be highly dependent on the presence of *KMT2D* mutations and raises the question as to whether KDM5i may be effective in other malignancies harbouring *KMT2D* lesions, or indeed mutations in other *KMT2* methyltransferases.

## Supporting information

Supplementary Figures, Tables, Methods and References

## Acknowledgements

We would like to thank EpiTherapeutics and Gilead for providing us with KDM5-inh1 and general advice throughout the project, in particular Lars-Ole Gerlach, Kristian Helin, Daniela Kleine-Kohlbrecher and Peter Staller (EpiTherapeutics).

## Authorship contributions

J.F. and J.H. conceived the study; J.H., G.P. A.M. and J.F. designed the study; J.H., A.D., G.P. and J.F. wrote the manuscript; J.G., P.J., J.O., S.I. and A.C. identified, contributed and prepared patient samples for the project; J.H., F.B.C and J.W. performed bioinformatic analysis; J.H., L.K., A.D., A.Y., T.R., A.F.A., S.A., K.C., M.P., J.D., D.B., K.K. and E.K. performed experiments; J.H., L.K., A.D., A.Y. and J.F. analysed the data; R.N., U.O. and A.M. contributed reagents and interpretation of data; All authors read, critically reviewed and approved the manuscript.

## Methods

### Cell culture

All cell lines were cultured in a 37°C, 5% CO_2_ humidified incubator using RPMI-1640 supplemented with 10% FBS, 1% L-glutamine and 1% Pen-Strep, except OCI-LY-1 and OCI-LY-7, which were cultured in IMDM with 20% FBS, 1% L-glutamine and 1% Pen-Strep (Supplementary Table 8). Cell lines were acquired from DSMZ or an institute tissue-bank. Identity was confirmed by STR sequencing and regularly checked by Sanger sequencing of unique mutations and for Mycoplasma contamination.

Primary FL cell-suspensions were defrosted at 37°C and layered onto 3ml of lymphoprep (STEMcell Technologies). Lymphocytes were isolated by centrifugation at 1150g for 12 minutes, washed in RPMI and resuspended in fresh RPMI before treatment. Written consent was obtained for the collection and use of specimens for research purposes with ethical approval obtained from the London Research Ethics Committee of the East London and the City Health authority (10/H0704/65, 06/Q0605/69) and Southampton and South West Hampshire (t228/02/t).

### Western blots

To assess histone mark levels, an isotonic lysis buffer (20mM Tris, 100mM NaCl, 5mM MgCl_2_, 10% glycerol, 0.2% NP40, 0.5mM DTT) and centrifugation was used to isolate nuclei, which where lysed in a high-salt buffer (50mM Tris, 600mM NaCl, 10% glycerol, 0.2% NP40, 0.5mM DTT) followed by sonication to fragment chromatin (Diagenode Bioruptor). Buffers were supplemented with phosphatase and Complete ULTRA protease inhibitor cocktails (Roche) and lysates quantified by Pierce 660nm Protein Assay Reagent (ThermoFisher). 1-2.5µg of nuclear protein was loaded in 4-12% Bis-Tris gels (NuPAGE), resolved by SDS-PAGE and transferred to polyvinylidene fluoride membranes via the iBlot™transfer device (Invitrogen). After blocking with 5% BSA/Milk in Tris-buffered saline, membranes were probed with primary antibody, stained with horseradish peroxidase-conjugated secondary antibodies (DAKO) and bands detected using ECL Plus (GE Healthcare) or SuperSignal West Femto (ThermoFisher). To examine multiple histone marks, equal amounts of protein were loaded onto multiple gels from a single loading solution, and quantified relative to H3.

For total protein analysis, western blots were performed as before except that cells were lysed in RIPA buffer (150mM NaCl, 1% NP-40, 0.5% sodium deoxycholate, 0.1% sodium-dodecyl-sulphate, 50mM tris(hydroxymethyl)aminomethane hydrochloride, pH 8.0), supplemented with 1X protease inhibitor and phosphatase inhibitor cocktail 2 and 3 (Sigma-Aldrich), for 30 minutes. Densitometry analysis was performed using ImageJ software and all the antibodies used are listed in Supplementary Table 9

### Proliferation and apoptosis assays

2000 cells were seeded in 100μl growth media in triplicate in 96-well plates 24h before treatment, followed by treatment with DMSO and 6 concentrations of KDM5i diluted 8-fold (0.0003-10μM) in 100μl of growth medium. The plates were incubated for five days, when viable cell numbers were determined using the Guava ViaCount assay (Millipore) or CellTitreGlo (Promega). The percentage of apoptotic cells was quantified by the Guava Nexin assay (Millipore), which measures binding of Annexin V to phosphatidyl serines and the incorporation of 7-AAD, a cell impermeable dye. Apoptotic cells were defined as Annexin V+/7-AAD+, early-apoptotic as Annexin V+/7-AAD- and nucleated debris as Annexin V-/7-AAD+.

For 10-day treatments, 20,000 cells were seeded in 1ml in 12-well plates and incubated overnight. The cells were then treated with DMSO or KDM5-inh1 (0.0024-10μM) and incubated for 5 days. Viable cell numbers were determined, and the cells re-seeded in triplicate in 96-well plates at 4000 cells/well and treated with the same concentration of KDM5-inh1 or DMSO as before. Viable cell numbers were determined by Guava ViaCount assay after a further 5-day incubation.

### Cell-cycle analysis

After treatment with DMSO or KDM5-inh1 for 72h, cells were permeabilized in ice-cold 70% ethanol, stored at -80°C and stained in PBS containing 50μg/ml propidium iodide and 100μg/ml RNase A. DNA content was then quantified using the YG610/20 filter on a Fortessa II flow cytometer.

### Surface IgM analysis

Cells were washed, re-suspended in FACS buffer (1% BSA, 4mM ethylenediaminetetraacetic acid (EDTA) and 0.15mM NaN_3_ in PBS) and stained for surface IgM expression (5×10^5^ cells/100µl) in the dark on ice for 30 minutes, using R-Phycoerythrin-conjugated anti-IgM (DAKO). Following incubation, cells were washed, re-suspended in FACS buffer and 1×10^4^ lymphocytes were acquired on a FACS Canto (BD Biosciences). Analysis of mean fluorescence intensities was performed with FlowJo software.

### Synergy analysis

Cells were treated with 5 concentrations of KDM5-inh1 for 5 days as per the standard proliferation assay described above, except that cells were also treated with 5 concentrations of S63845 and Venetoclax for 2 days or Ibrutinib for 3 days. Synergy was assessed for each combination using the DrugComb portal (https://drugcomb.fimm.fi/analysis/).

### RNA extraction

RNA was extracted using QIAGEN RNeasy kits including an on-column DNase step. RNA for sequencing was determined to be of high-quality by Agilent Bioanalyser or Tapestation (RIN > 9.5).

### cDNA synthesis and qRT-PCR

cDNA was synthesised using the high capacity cDNA reverse transcription kit (ThermoFisher) and qPCR performed using the SsoAdvanced(tm) Universal SYBR^®^ Green Supermix (BioRad). Reactions were performed in triplicate and normalised to GAPDH. All primer sequences are listed in Supplementary Table 10.

### ChIP-PCR and –seq

ChIP reactions were prepared using a modified version of the Active Motif ChIP-IT High Sensitivity Kit. For cell lines, 5-15 million cells were treated for 72h with DMSO or 1μM KDM5-inh1, cross-linked in a 1% formaldehyde/PBS solution for 5 minutes, washed in PBS, and nuclei isolated by 5 minute incubation on ice in a cytoplasmic lysis buffer (50mM Tris·Cl, 140mM NaCl, 1.5mM MgCl_2_, 0.5% (v/v) Nonidet P-40 (1.06g/ml)) and centrifugation. Nuclei were sonicated using a BioRuptor for 10-20 cycles of 30s on (high)/60s off. After confirming correct fragmentation of input DNA, ChIP reactions were performed overnight at 4°C (antibodies listed in Supplementary Table 9), followed by DNA precipitation with agarose beads, reversal of cross-links and purification of DNA by columns. Samples were analyzed by qPCR using the probes listed in Supplementary Table 10. Lymphocytes from primary FL cell-suspensions were examined identically except that 1-5 million cells were used per ChIP and that the cells were treated for 48h. For ChIP-seq, the chromatin was spiked with 15ng of Drosophila chromatin (Active Motif; 53083) and ChIP was performed with an anti-*Drosophila* chromatin antibody (Active Motif; 61686) alongside the H3K4me3/H3K4me1 antibodies.

Libraries were prepared for sequencing using the NEBNext Ultra II and Multiplex Oligios for Illumina kits (New England Biolabs) according to the manufacturers protocol. Briefly, ChIP and input DNA were end-repaired, adaptors ligated and size-selected using SPRIselect beads for 300-400bp DNA fragments and amplified by PCR. Correct library size (400-500bp) was confirmed by Tapestation. Sequencing was performed on the Illumina HiSeq 4000 to generate 75bp paired-end reads or NextSeq 500 to generate 40bp single-ended reads.

### CRISPR

To generate *KMT2D* mutant cells, four pooled guide-RNAs (gRNAs) were designed targeting exon 3 of *KMT2D*, whilst individual gRNAs were used for KDM5A/KDM5C (Supplementary Table 11). gRNAs were combined with tracrRNA at equimolar concentrations, heated at 95°C before cooling to anneal. 460pmol of the gRNA/tracrRNA pool was then complexed with 401pmol of Cas9 protein (Alt-R Cas9 Nuclease 3NLS; IDT) and transfected by Nucleofection (Supplementary Table 12). After transfection, cells were left to recover in 4ml of complete growth medium and after 48h a cell-sorter was used to isolate single cell clones. Editing was identified by Sanger sequencing and validated by TA-cloning for complex mutations (Supplementary Table 1). To correct the 1bp insertion present in *KMT2D* within SU-DHL-8 cells, CRISPR was performed as above except that a donor-template containing 119bp of WT sequence, with a silent mutation to alter the PAM site, was co-transfected alongside the gRNA (targeting the mutation site) and Cas9 protein.

### Xenograft studies

SU-DHL-6 xenograft studies were performed by Crown Bioscience Inc. (Beijing). 24h after irradiation with Co^60^ (150 rads), 5×10^6^ SU-DHL-6 cells (in 0.1ml PBS mixed with matrigel 1:1) were inoculated subcutaneously into the right flank of NOD/SCID mice (weighing 18-20g). Once tumours reached an average size of 100mm^3^, mice were randomized into three groups of 10; vehicle (6% Captisol + 94% ddWater, pH=2), KDM5-inh1 and ibrutinib. Mice were orally dosed daily with 50mg/kg KDM5-inh1 and 10mg/kg ibrutinib up to 21 days, with a scheduled dosing holiday for the KDM5-inh1 group between days 8-14. Six mice were additionally randomized into two groups (n=3) and treated with vehicle or 50mg/kg KDM5-inh1 for 1 week. Tumour volumes were calculated in two dimensions 3x a week. Mice were euthanized when the mean tumour size of the vehicle group exceeded 2000mm^3^ or once the study endpoint was reached.

