## Supplementary Figures, Tables, Methods and References for "KDM5 inhibition offers a novel therapeutic strategy for the treatment of *KMT2D* mutant lymphomas"

### Supplementary Information

#### Supplementary Methods:

##### Mutational profiling of cell lines

Targeted resequencing was performed on 28 frequently mutated genes using the Access Array platform (Fluidigm) as previously described (1). Amplification was performed in multiplex using 50ng genomic DNA whilst sequencing was performed on pooled libraries using the Illumina MiSeq platform. Data were aligned to hg19 using Bowtie2 (59) and VarScan2 (60) was used to call variants, including SNPs and short indels, implementing a threshold VAF > 5%, a read depth > 10 and a Fisher's exact test *P* value of <0.05. Identified variants were annotated using SNPnexus. For OCI-LY-18, whole exome capture libraries were prepared using the Agilent SureSelect Human All Exon Kit V5 (Agilent Technologies). Paired-end multiplex sequencing of samples was performed on the Illumina HiSeq 2500. Data were aligned to hg19 using the Burrows-Wheeler Aligner (BWA) (61), and converted into BAM files with the exclusion of PCR duplicates and removal of low quality reads using Picard (version 1.86), whilst the Genome Analysis Toolkit (GATK) version 2.3.9 (62) was used for realignment of discordantly mapped read pairs around indels and recalibration of base quality scores. Somatic single nucleotide variants (SNVs) were identified using the Strelka pipeline and independently verified by Sanger sequencing (Supplementary Table 1).

##### RNA-seq and analysis

mRNA isolation and strand-specific cDNA libraries were prepared using the Illumina TruSeq stranded mRNA Prep kit according to the manufacturer's instructions. For SU-DHL-6 and OCI-LY-18, two hundred cycles of sequencing on the Illumina HiSeq 2500 instrument were performed to generate 2x100bp paired-end sequencing reads. HT libraries were sequenced by 300 cycles of sequencing on the HiSeq 4000 to generate 2x150bp paired-end sequencing reads while WSU-DLCL2/WSU#22 cells were sequenced using 2x40bp reads generated on the NextSeq 500. Raw data was aligned to hg19 using STAMPY (63) and BWA (61). The number of uniquely aligned reads (quality score $q$ > 10) aligned to the exonic region of each gene were counted using HTSeq (64) based on the Ensembl annotation (version 75). Only genes that achieved at least one read per million reads (CPM) in at least 3 samples were kept. Aligned reads numbers were further normalised using the 'cqn' method (65), accounting for gene length and GC content. Differential expression (DE) analysis was then performed using the edgeR R package (66), employing the generalised linear model (GLM) approach, for the treated versus control pairwise matched comparisons. DE genes were selected based on a false discovery rate (FDR) of <0.05 and a fold change >2 or <0.5.

Raw read counts across 55,765 annotated genes for 97 FL and 75 DLBCL samples were obtained from the ICGC and converted to RPKM expression values. Gene-level transcription estimates of 20,531 genes (log2(transformed RSEM normalized count + 1) (level_3 data)) from were obtained from the TCGA Large B-cell Lymphoma (DLBC) cohort (n=48) using the UCSC Xena Browser (https://xenabrowser.net/datapages/). RPKM expression values from healthy GC B-cells were downloaded from the BLUEPRINT EPICO data portal (http://blueprint-data.bsc.es/release_2016-08/#!/).

To identify clusters within the KDM5-inhibition/*KMT2D* mutation RNA-seq datasets, log2 fold change values from 897 genes DE following either KDM5-inhibition or

*KMT2D* mutation were selected. K-means clustering was performed for 1-20 clusters using the kmeans function in R, with six identified as the optimal cluster number by plotting the cluster number verses the “within-cluster sum of squares”.

**ChIP-seq analysis**

To normalize ChIP-seq data, protocols published by Active Motif and used to investigate other epigenetic drugs were followed (67). Input FASTQ files were mapped to hg19 using Bowtie 2 (59), whilst H3K4me3/me1 FASTQ files were mapped to hg19 and dm3. The number of reads mapping to the dm3 genome, derived from the spike-in, were used to calculate a scaling factor and sequencing reads were randomly removed from samples requiring normalization using SAMtools (68) (samtools view -b -s <scaling.factor> <filename.bam> > <scaled.bam>) i.e. samples with higher spike-in read counts were downscaled so that the overall read number would be relative to the level of the histone mark in that sample. Peaks were identified using the broad source option in MACS2 (macs2 callpeak -t <filename.bam> -c <input.bam> --broad -n <filename> -g hs --broad-cutoff 0.1 -q 0.05) and regions known to result in poor quality sequencing (https://www.encodeproject.org/annotations/ENCSR636HFF/) removed using BEDtools (69). Analysis of differential H3K4 methylation was performed in R using EdgeR within the DiffBind package (70). Peak annotation was performed using ChIPseeker package (71) and Genomic Regions Enrichment of Annotations Tool (GREAT) (72). The overlap with publicly available ChIP-seq datasets was performed using Intervene (73) to generate pairwise intersection matrices, BEDtools intersect (69) to assess individual relationships and the Genomic Association Tester to calculate significance (74).

To examine the relationship between H3K4me3 and H3K4me1 at promoters following KDM5-inhibition, 1kb intervals were generated from H3K4me3 positive promoters, centered (+/- 500bp) on the TSS. Bedtools coverage (bedtools coverage -a <Intervals.bed> -sorted -b <chip.bam>) was then used to count reads in each interval and calculate fold changes following KDM5-inh1 exposure. Profile plots for histone marks were generated using the computeMatrix and plotProfile tools from Deeptools (58).

To assess the enrichment of histone marks and epigenetic regulators at promoters (TSS +/- 500bp), TSS locations for each gene were downloaded from BioMart and any overlapping intervals merged. Intervals overlapping genes in each cluster were then selected and Deeptools MultiBigwig summary (MultiBigwigSummary bins -b <bigwig files> --bed <Cluster.bed> --outRawCounts <filename> ) (58) used to calculate a summary score for each promoter using bigwig files downloaded from the ENCODE or BLUEPRINT projects. The control promoters were generated as above, using genes that were not DE or differentially methylated in any RNA- or ChIP-seq experiment, and then overlapped with H3K27me3 deposition (31) to identify non-bivalent (H3K4me3+/H3K27me3-) and bivalent (H3K4me3/H3K27me3+) promoters. Mean values were then scaled between clusters for each dataset using the scale function in R.

All generated ChIP-seq data have been deposited in the GEO database under the accession number GSE114492 and GSE152642.

**Pathway enrichment**

Pathway enrichment on genes proximal to ChIP-seq intervals was performed using GREAT (72). For RNA-seq data and analysis of promoters with differential H3K4me3 deposition, Gene Set Enrichment Analysis (GSEA) was performed (33) using pre-ranked option and "classic" scoring. To interrogate publicly available datasets from different B-cell/lymphoma studies, we used a manually curated version of the lymphochip database (28) where key studies from the literature were manually added.

#### Supplementary Figures:

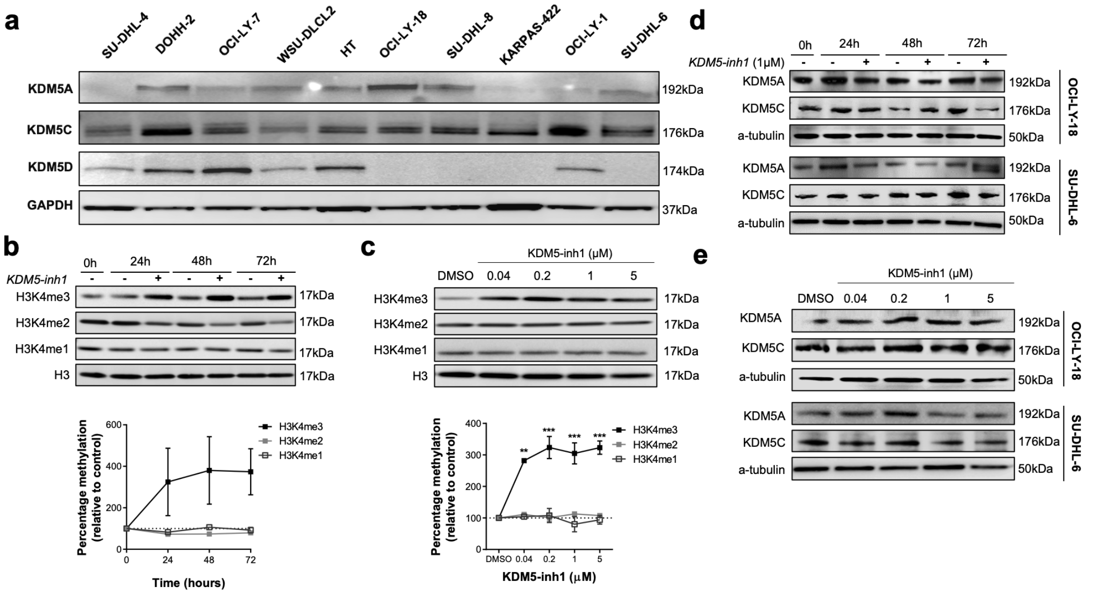


**Supplementary Figure 1. Validation of KDM5 expression and effect of KDM5-inhibition on H3K4me in DLBCL cell lines.** **(a)** Expression of KDM5A, C and D was confirmed by western blot in 10 DLBCL cell lines. OCI-LY-18 cells were **(b)** treated with DMSO or 1μM KDM5-inh1 for increasing lengths of time and **(c)** DMSO or increasing concentrations of KDM5-inh1 for 48h, followed by western blots to quantify H3K4me3/me2/me1 and H3 levels. Upper panels display representative western blots and lower panels display the quantified western blots relative to H3. Data are the mean ± SEM of 3 independent experiments. Statistical significance was determined using a one-way ANOVA with a Dunnett's post-test versus untreated control, where ** $P<0.01$ and *** $P<0.001$. KDM5A and KDMC5 expression was examined in OCI-LY-18 (upper panels) and SU-DHL-6 (lower panels) cells treated with **(d)** DMSO or 1μM KDM5-inh1 for increasing lengths of time and **(e)** DMSO or increasing concentrations of KDM5-inh1 for 48h**.**

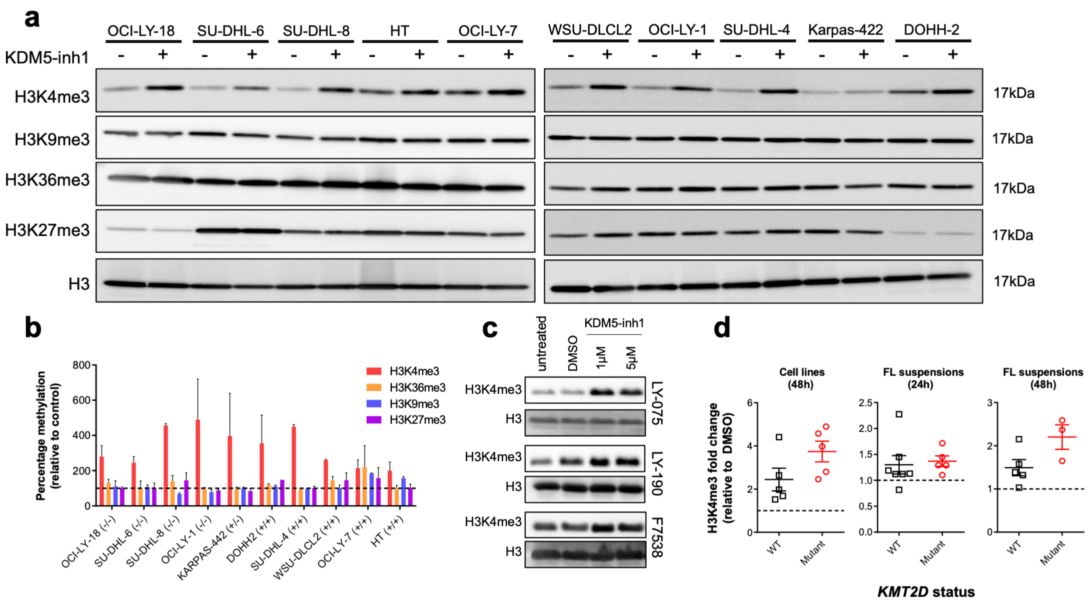


**Supplementary Figure 2. KDM5 inhibition increases H3K4me3 but not H3K9me3, H3K36me3 or H3K27me3.** 10 DLBCL cell lines were treated with DMSO or 1µM KDM5-inh1 for 48h, followed by western blots to quantify H3K4me3, H3K9me3, H3K36me3 and H3K27me3 levels. Representative western blots are displayed in **(a)** and the quantification of histone marks relative to H3 in **(b)**. Data are the mean ± SEM of 2 independent experiments. **(c)** H3K4me3 levels and PARP cleavage (no difference – data not shown) were analysed by western blot in primary FL cell suspensions exposed to 1µM and 5µM KDM5-inh1 for 24 and 48h. H3K4me3 levels after 48h are displayed in **(c)** and the quantification of H3K4me3 relative to H3, in KMT2D mutant and WT samples exposed to 1µM KDM5-inh1, in **(d)**. The quantification of mutant and WT cell lines after exposure to 1µM KDM5-inh1 for 48h is also displayed. The primary cell analysis was performed on 5 mutant *KMT2D* and 7 WT cell suspensions at 24h, and 3 mutant and 5 WT at 48h.

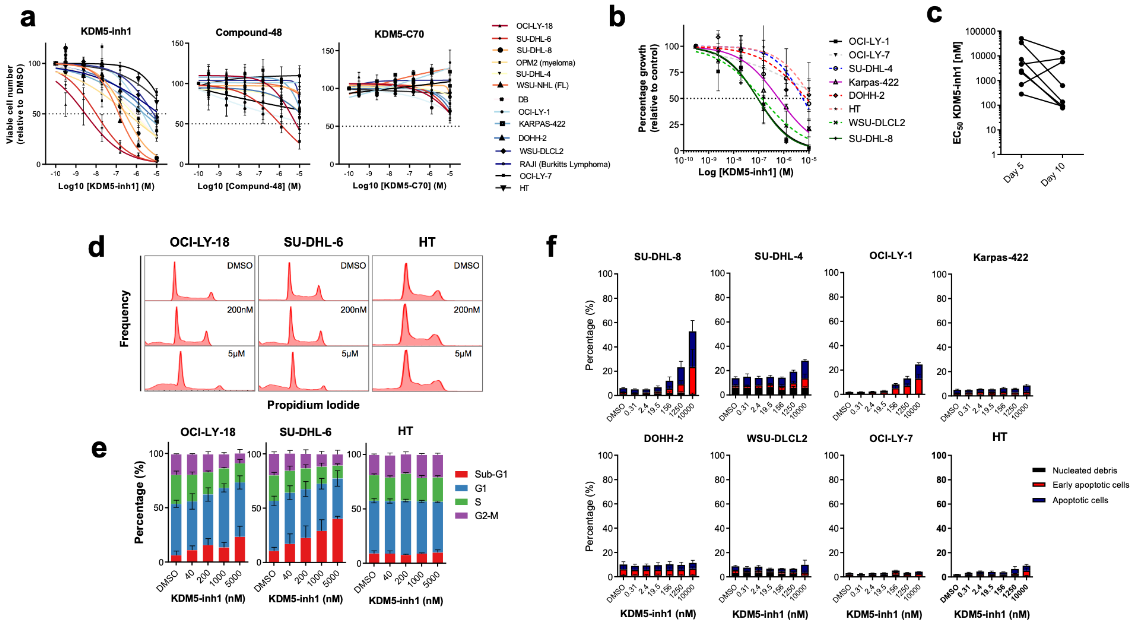


**Supplementary Figure 3. KDM5-inhibition is cytostatic and cytotoxic in cell lines. (a)** DLBCL, FL, myeloma and Burkitt's lymphoma cell lines were treated with DMSO or increasing concentrations of KDM5-inh1, Compound-48 or KDM5-C70, and viable cells quantified after 5 days. Data are the mean ± SEM of 3-6 independent experiments. **(b)** 8 DLBCL cell lines were treated with DMSO or increasing concentrations of KDM5-inh1 for 10 days, with the cells re-seeded in fresh drug/media after 5 days. Data are the mean ± SEM of 3 independent experiments. **(c)** Mean $EC_{50}$ values after 5 and 10 days. **(d+e)** DNA content was quantified in 2 sensitive (OCI-LY-18 and SU-DHL-6) and an insensitive (HT) DLBCL cell line using propidium-iodide staining and flow cytometry after treatment with increasing concentrations of KDM5-inh1 for 72h. Representative histograms are shown in **(d)** and quantification of cell cycle phases in **(e)**. **(f)** Induction of apoptosis was quantified by the Guava Nexin assay in 8 DLBCL cell lines treated with DMSO or increasing concentrations of KDM5-inh1 for 5 days**.** Data are the mean ± SEM of 3 independent experiments.

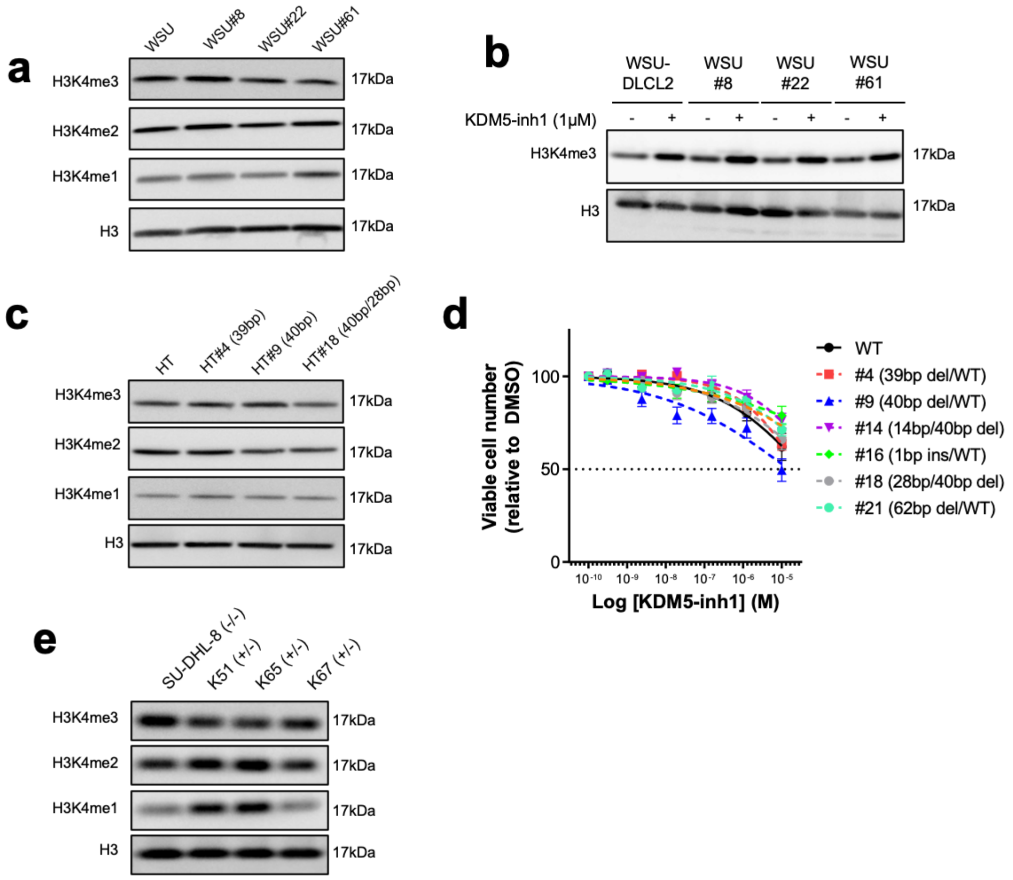


**Supplementary Figure 4. Histone mark levels in *KMT2D* CRISPR models. (a)** H3K4me3/me2/me1 and H3 levels were quantified by western blot in WSU-DLCL2 parental and *KMT2D* mutant cells. **(b)** H3K4me3 and H3 protein levels were quantified by western blot in WSU-DLCL2 parental and KMT2D mutant cells treated with DMSO or 1µM KDM5-inh1 for 48h. **(c)** H3K4me3/me2/me1 and H3 levels were quantified by western blot in parental HT and *KMT2D* mutant HT cells. **(d)** HT cells and 6 *KMT2D* mutant HT clones were treated with DMSO or increasing concentrations of KDM5-inh1 for 5 days, and then quantified by Guava ViaCount. The size of induced deletions is indicated in brackets. **(e)** H3K4me3/me2/me1 and H3 levels were quantified by western blot in homozygous mutant *KMT2D* SU-DHL-8 parental cells and in three clones with a corrected allele. Results are representative of 3 independent experiments.

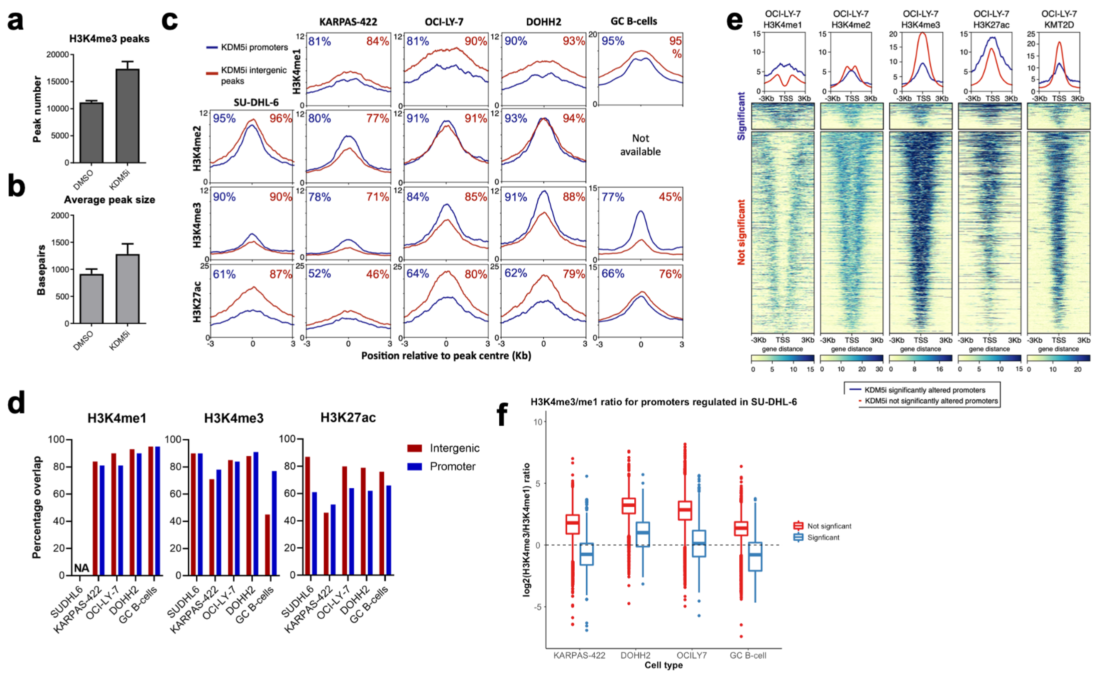


**Supplementary Figure 5. KDM5-inhibition targets regions marked by high H3K4me1. (a)** Mean number of H3K4me3 peaks and **(b)** mean peak width identified by ChIP-seq in DMSO and 1μM KDM5-inh1 treated SU-DHL-6 cells. **(c)** plotProfile (58) was used to compare basal levels of H3K4me3/H3K4me1 in lymphoma cell lines (ENCODE) and primary GC B-cells (BLUEPRINT) at promoters and intergenic regions displaying significantly altered levels of H3K4me3 following KDM5-inhibition. **(d)** Summary of percentage overlaps between KDM5-inhibition regulated intergenic regions or promoters, and H3K4me1, H3K4me3 and H3K27ac peaks. **(e)** Heatmaps of H3K4me3/me2/me1 levels (ENCODE) and KMT2D binding (10) in OCI-LY-7 cells, showing differences between promoters significantly altered (blue) or otherwise (red) in SU-DHL-6 cells treated with DMSO or 1μM KDM5-inh1 for 72h. **(f)** Deeptools (58) was used to calculate summary scores from H3K4me3 and H3K4me1 ChIP-seq datasets, using H3K4me3 peaks called in SU-DHL-6 cells. Peaks were subdivided on the basis of being significantly altered by KDM5-inhition in SU-DHL-6 cells, and log2 ratios of the H3K4me3 score/H3K4me1 score calculated for each examined cell type.

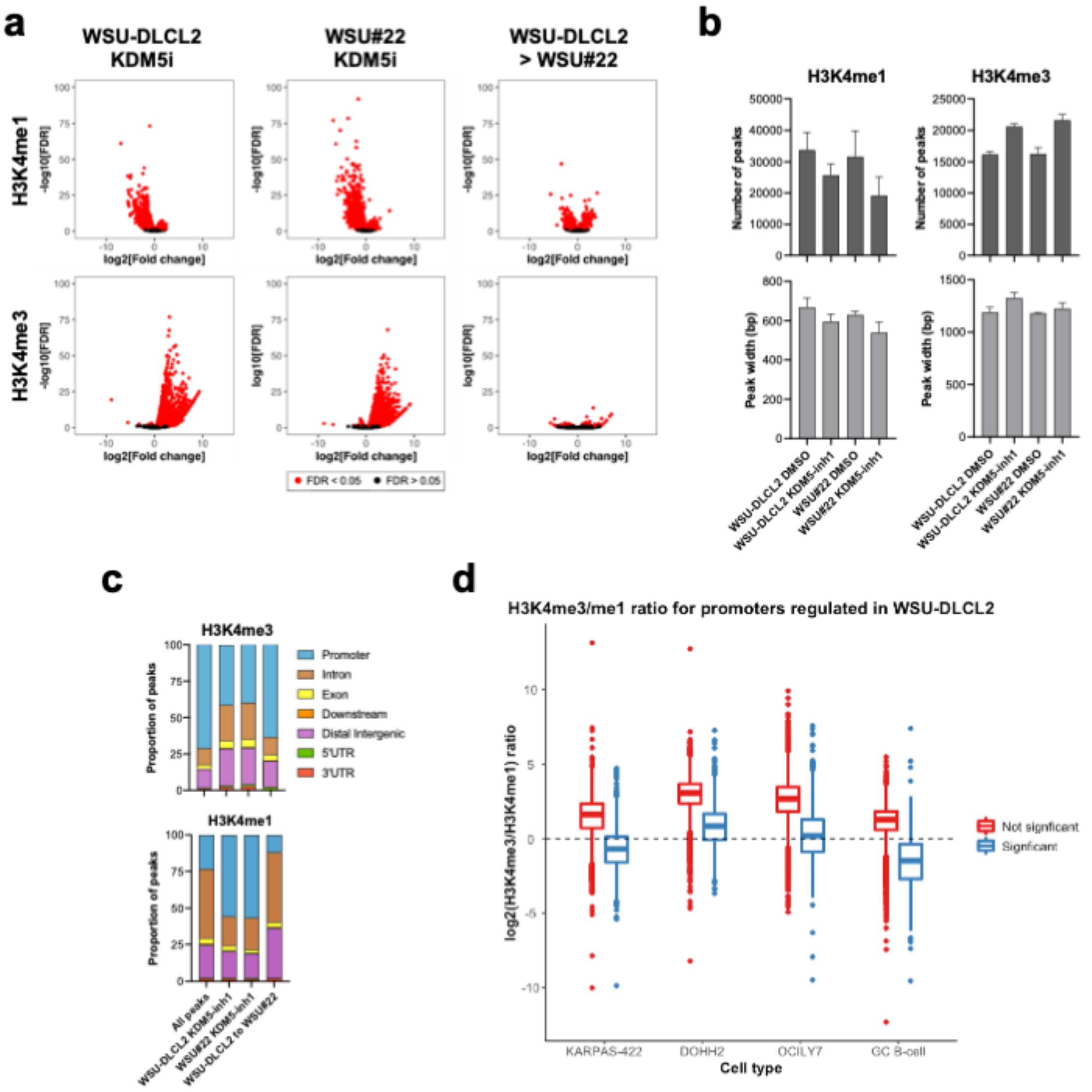


**Supplementary Figure 6. KDM5-inhibition reduces global levels of H3K4me1 whilst increasing H3K4me3. (a)** Volcano plots displaying changes in H3K4me3 or H3K4me1 following ChIP-seq analysis of 1μM KDM5-inh1 treated WSU-DLCL2 and WSU#22$^{-/+}$ cells, or between WSU-DLCL2 and WSU#22$^{-/+}$ cells. **(b)** Number and width of H3K4me1/H3K4me3 peaks in DMSO or 1μM KDM5-inh1 treated WSU-DLCL2 and WSU#22$^{-/+}$ cells. **(c)** Annotation of differential H3K4me1/H3K4me3 peaks following 1μM KDM5-inh1 treatment or between WSU-DLCL2 and WSU#22$^{-/+}$ cells, alongside total peak annotations. **(d)** Deeptools (58) was used to calculate summary scores from H3K4me3 and H3K4me1 ChIP-seq datasets, using H3K4me3 peaks called in WSU-DLCL2/WSU#22$^{-/+}$ cells. Peaks were subdivided on the basis of being significantly altered by KDM5-inhition in WSU-DLCL2/WSU#22$^{-/+}$cells, and log2 ratios of the H3K4me3 score/H3K4me1 score calculated for each examined cell type.

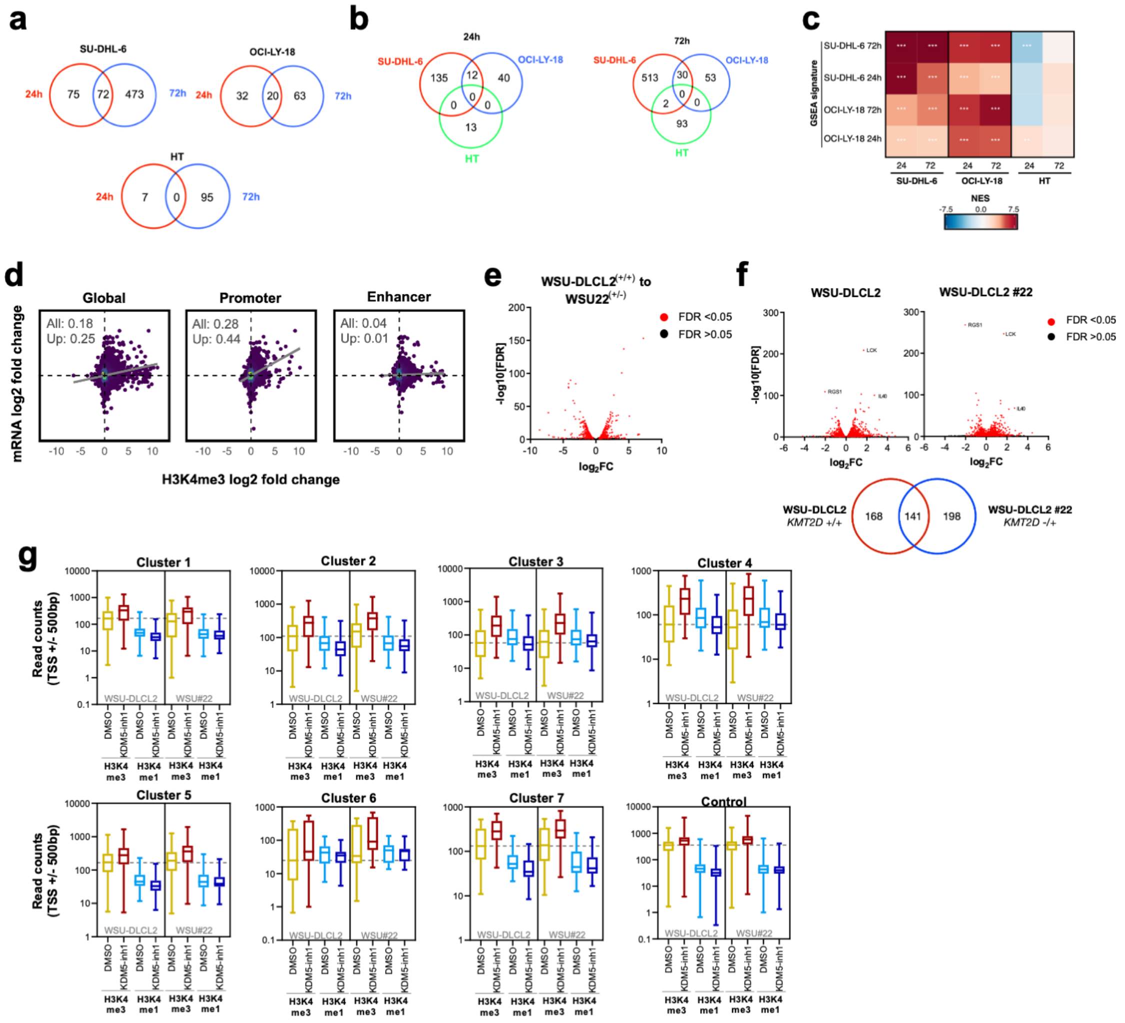


**Supplementary Figure 7. Epigenetic and transcriptomic analysis of KDM5-inhibititon.** Overlap between KDM5-inh1 regulated genes at **(a)** 24h and 72h in SU-DHL-6, OCI-LY-18 and HT cells or **(b)** between cell lines at 24h and 72h. **(c)** GSEA was performed using signatures derived from each cell line (y axis) to examine their enrichment in all cell types and timepoints (x axis). Note that signatures derived from HT cells were not enriched in any condition, likely due to low numbers of protein-coding genes. **(d)** Correlation analysis between mRNA expression and H3K4me3 levels following KDM5-inhibition for all H3K4me3 peaks (Global), promoters or putative enhancers. Peaks were annotated according to the nearest genes by ChIPseeker (71). Pearson's correlation coefficients are indicated in the top left of each panel for all genes and genes with a log2 fold change > 0 (Up). **(e)** Volcano plot showing DE genes between WSU-DLCL2 and WSU#22$^{-/+}$ cells, with significant genes highlighted in red. **(f)** Volcano plots showing DE genes induced following exposure of WSU-DLCL2 and WSU#22$^{-/+}$ cells to 1μM KDM5-inh1 for 72h, with significant genes highlighted in red. The overlap between the two is shown in the lower panel. **(g)** Boxplots showing H3K4me3/H3K4me1 promoter (TSS +/-500bp) read counts across ChIP-seq datasets, of genes from all identified clusters (Figure 3f) plus control promoters, which were selected on the basis of being H3K4me3+ in WSU-DLCL2/ WSU#22$^{-/+}$ cells but not altered at the mRNA or H3K4me1/H3K4me3 level in any of our analyses.

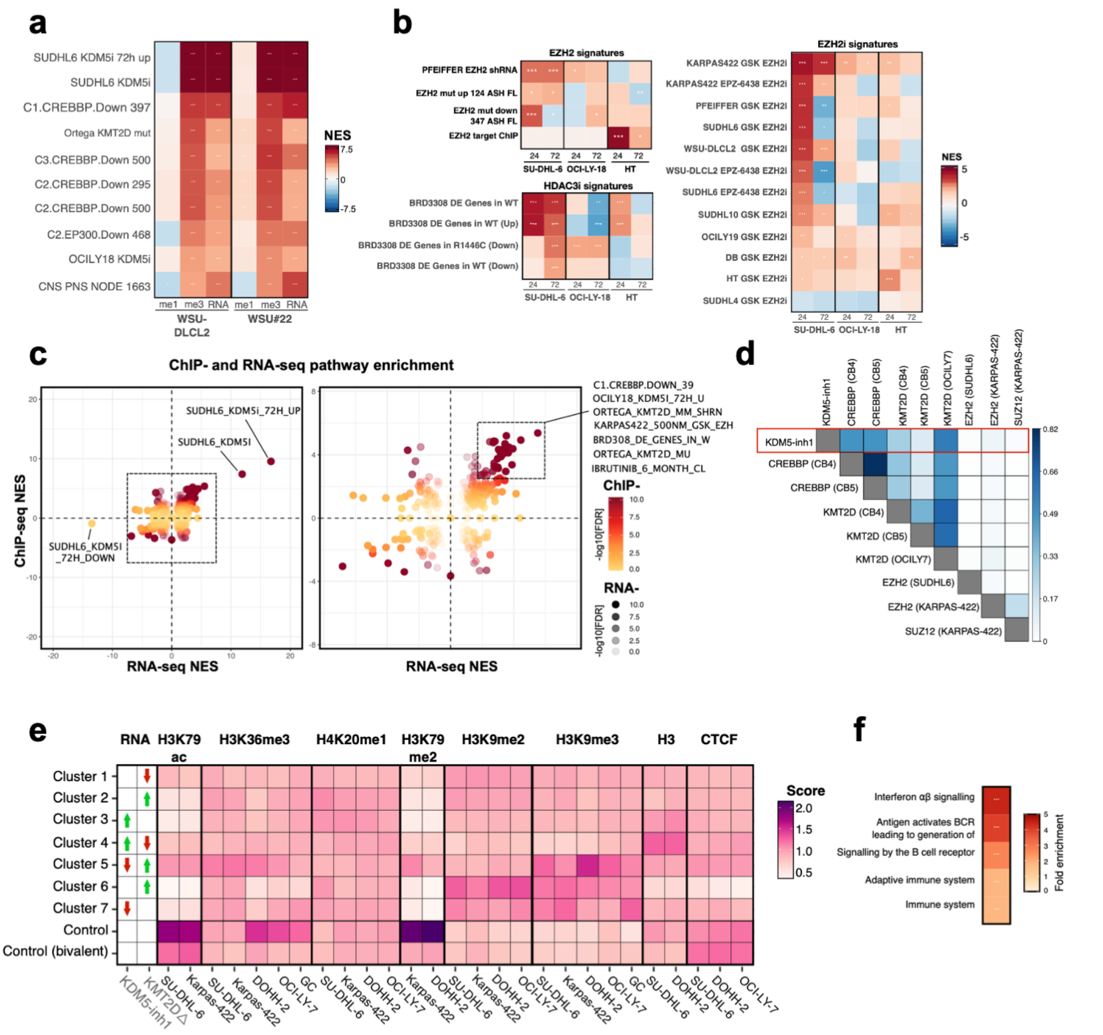


**Supplementary Figure 8. Pathway analysis of RNA- and ChIP-seq data**. **(a)** GSEA of RNA and H3K4me3/H3K4me1 (promoter) datasets following KDM5-inhibition, using a manually curated database of B-cell/lymphoma signatures. **(b)** GSEA of KDM5-inh1 RNA-seq data for EZH2, EZH2i (30,31) and HDAC3i (29) gene signatures. **(c)** Plot comparing GSEA of B-cell/lymphoma signatures from ChIP- and RNA-seq data in SU-DHL-6 cells at 72h. The left panel shows that genes upregulated at an mRNA level by KDM5-inhibition are also strongly positively enriched at the H3K4me3 level, in contrast to downregulated genes which show no H3K4me3 enrichment. The magnified right panel shows other notable datasets. **(d)** Intersection matrix (73) showing overlap between KDM5-inhibition regulated regions and KMT2D (9,10), CREBBP (36) and EZH2/SUZ12 (32) binding. **(e)** Deeptools (58) was used to calculate summary scores at the promoters (TSS+/-500bp) of genes in each cluster, plus non-bivalent (H3K4me3+/H3K27me3-) and bivalent (H3K4me3/H3K27me3+) control promoters, for ChIP-seq datasets of histone mark deposition and CTCF binding (ENCODE/BLUEPRINT). The overall direction of change in RNA expression, following KDM5i or *KMT2D* loss (Figure 3f), is indicated for each cluster in the first two columns. **(f)** Plot showing enrichment of Reactome signatures, calculated using GREAT, in H3K4me3 ChIP-seq data following exposure of SU-DHL-6 cells to 1μM KDM5-inh1 for 72h.

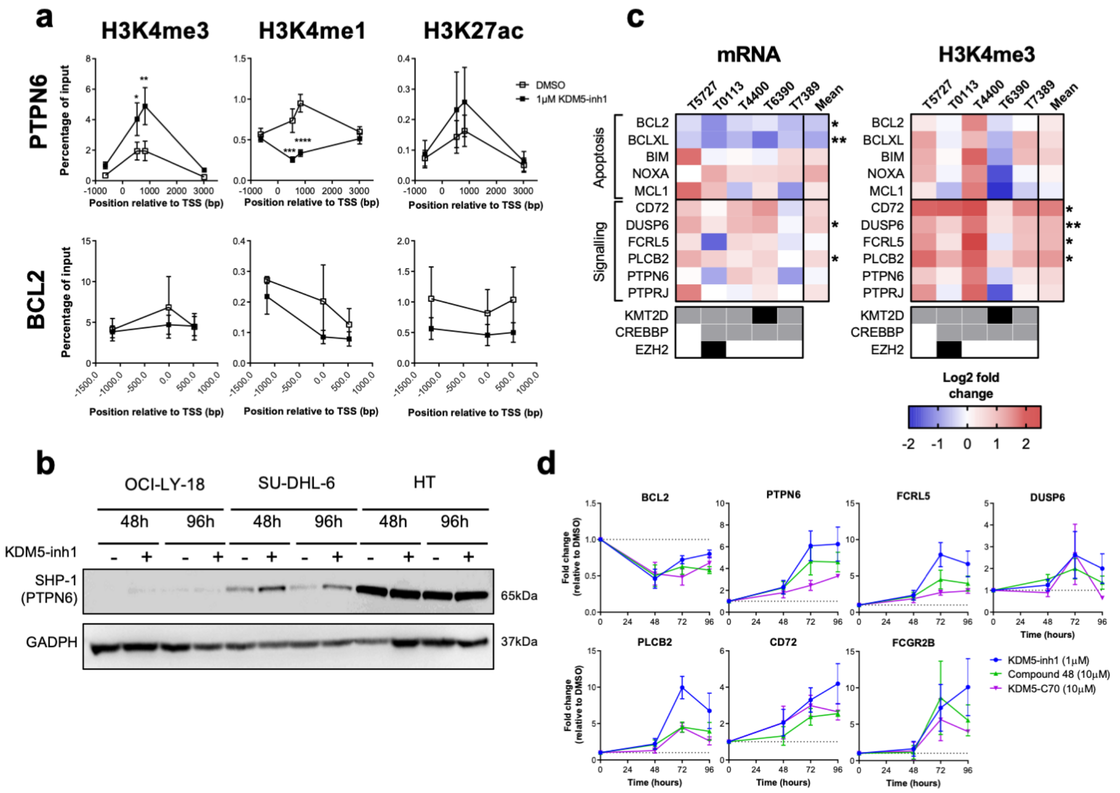


**Supplementary Figure 9. KDM5-inhibiton induces BCR-signalling regulators (a)** ChIP-PCR analysis of H3K4me3, H3K4me1 and H3K27ac at the *PTPN6* and *BCL2* promoters, following 72h DMSO or 1μM KDM5-inh1 exposure in SU-DHL-6 cells. Data was expressed as the percentage of input and plotted relative to the Transcription Start Site (TSS) of *PTPN6/BCL2.* Data are the mean ± SEM of 3 independent experiments. Statistical significance was calculated using a two-way ANOVA with a Dunnett's post-test, where * P <0.05, ** P<0.01, *** P<0.001 and **** P<0.0001. **(b)** Western blots of SHP-1 protein in OCI-LY-18, SU-DHL-6 and HT cells treated with DMSO or 1μM KDM5-inh1 for 48h and 96h. **(c)** Five primary FL cell suspensions were exposed to DMSO or 1μM KDM5-inh1 for 48h, followed by qRT-PCR and H3K4me3 ChIP-PCR analysis of KMD5i target genes. A summary heatmap is displayed, with the mutational status of patient samples indicated by grey (mono-allelic) and black (bi-allelic) squares in the lower panel. Statistical significance was calculated using a paired students T-test, where * P <0.05 and ** P <0.01. **(d)** qRT-PCR analysis of KDM5-inhibition target genes was performed on SU-DHL-6 cells treated with 1μM KDM5-inh1 or 10μM Compound 48 and Compound 70 for 48h, 72h and 96h. Data are the mean ± SEM of 3 independent experiments.

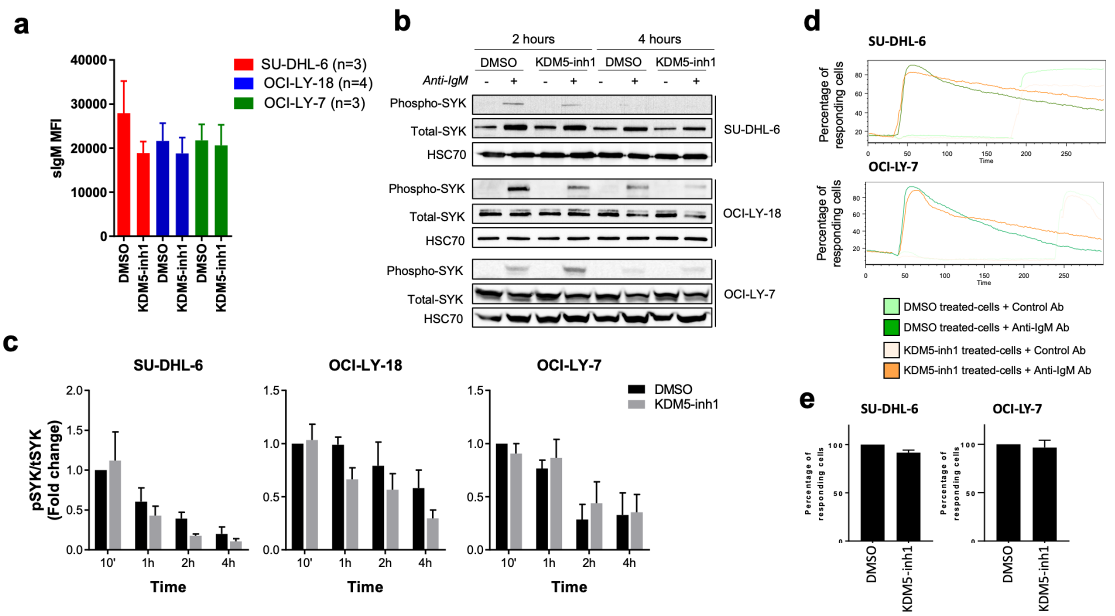


**Supplementary Figure 10. KDM5-inhibtion exposure results in diminished BCR signalling (a)** Surface IgM (sIgM) expression was quantified by flow cytometry in SU-DHL-6, OCI-LY-18 and OCI-LY-7 cells exposed to DMSO or 1µM KDM5-inh1 for 72h. **(b+c)** Activation of the BCR-associated kinase SYK was investigated in SU-DHL-6, OCI-LY-18 and OCI-LY-7 cells pre-treated with DMSO or 1µM KDM5-inh1 for 72h, followed by stimulation with anti-IgM F(ab')$_2$ antibody for 2h and 4h. Representative western blots for phospho-SYK, total SYK, and HSC70 are displayed in **(b)**, with quantification displayed in **(c)**. Data are the mean ± SEM of 3 independent experiments. SU-DHL-6 and OCI-LY-7 cells were treated with either DMSO or 1µM KDM5-inh1, and labelled with the calcium sensitive dye Fluo-3-AM and analysed by flow cytometry before and after addition of either F(ab')$_2$ anti-IgM or control antibody. **(d)** Panels showing sIgM-mediated intracellular calcium mobilisation as a percentage of responding cells, with quantification displayed in **(e)**. Data are the mean ± SEM of 2 independent experiments.

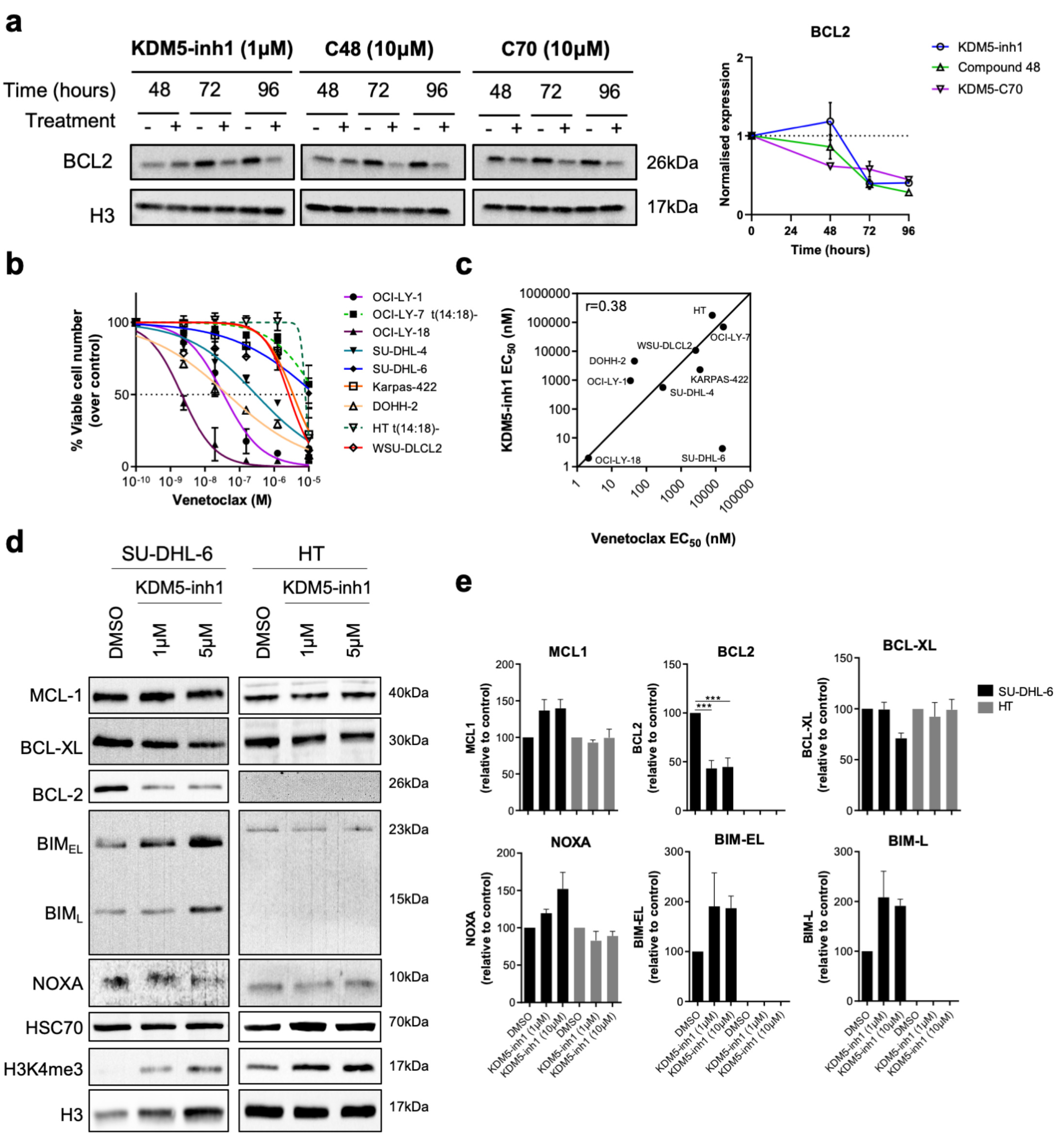


**Supplementary Figure 11. Response to BCL2i and expression analysis of BCL2 family members (a)** SU-DHL-6 cells were treated with 1μM KDM5-inh1 or 10μM Compound-48 or KDM5-C70 for 48h, 72h and 96h, followed by western blot analysis of BCL2 expression relative to H3. The quantification from three independent western blots (mean ± SEM) is shown in the right panel. Note that the H3 loading control was from the same experiment in Figure 1c. **(b)** Viable cells counts following exposure to DMSO or increasing concentrations of Ventoclax for 48h. t(14;18)- cell lines are indicated with dashed lines. **(c)** Correlation of Venetoclax and KDM5-inh1 $EC_{50}$ values, with the Pearson's correlation coefficient indicated. Results are representative of 3 independent experiments. **(d)** The expression of BCL2 family members was investigated by western blot analysis of SU-DHL-6 cells and HT cells treated with DMSO or 1μM and 5μM KDM5-inh1 for 2 days, with HSC70 used as a loading control. Representative western blots of 3 independent experiments are shown in **(d)**, with the quantification relative to HSC70 for in **(e)**. Statistical significance was determined using a one-way ANOVA with a Dunnett's post-test versus untreated control, where *** P < 0.001.

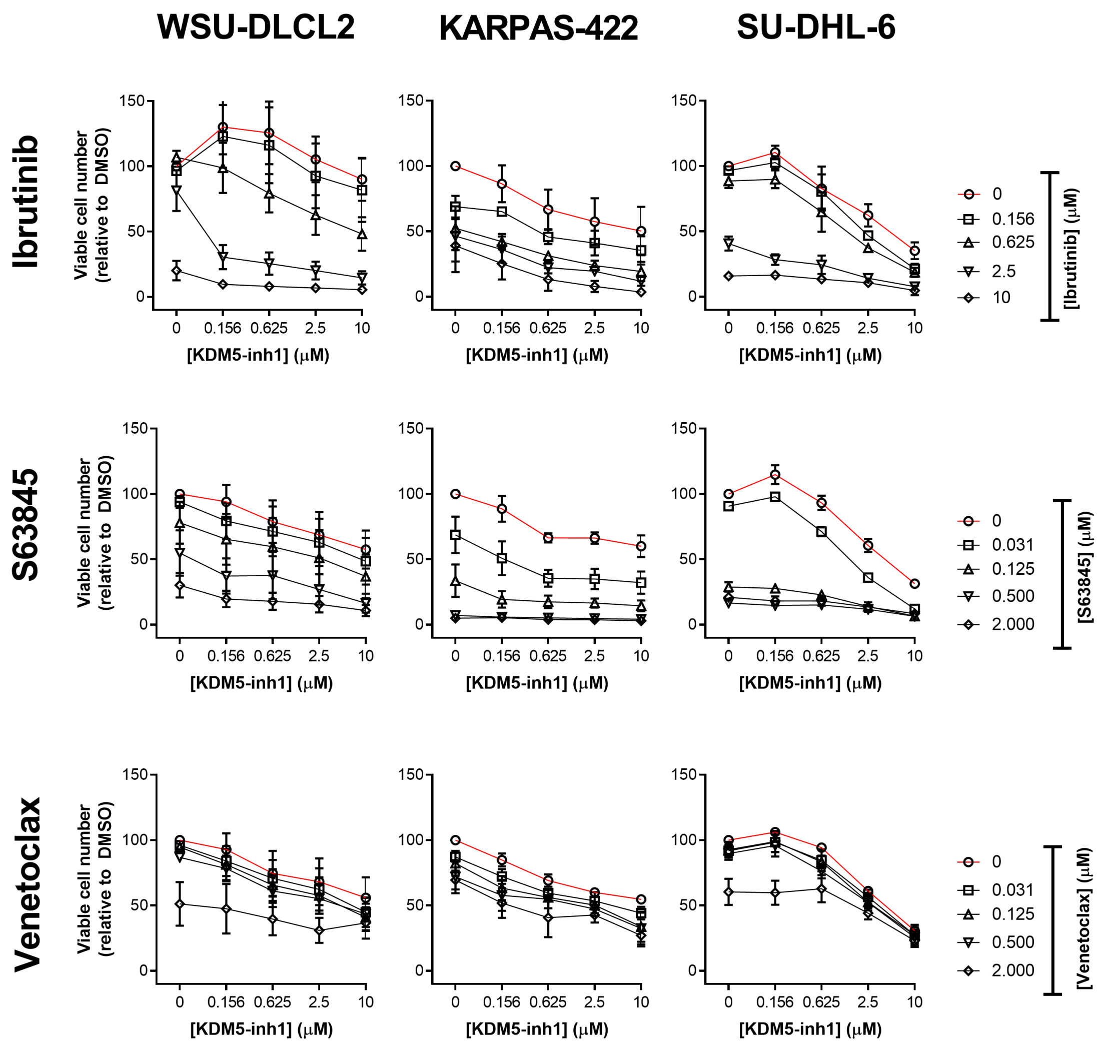


**Supplementary Figure 12. KDM5-inh1 synergises with BTKi and MCL1i.** WSU-DLCL2, KARPAS-422 and SU-DHL-6 cells were exposed to DMSO or increasing concentrations of KDM5-inh1 for 5 days, and combined with DMSO or increasing concentrations of Venetoclax or S63845 for 2 days or Ibrutinib for 3 days. Data are the mean ± SEM of 3 independent experiments.

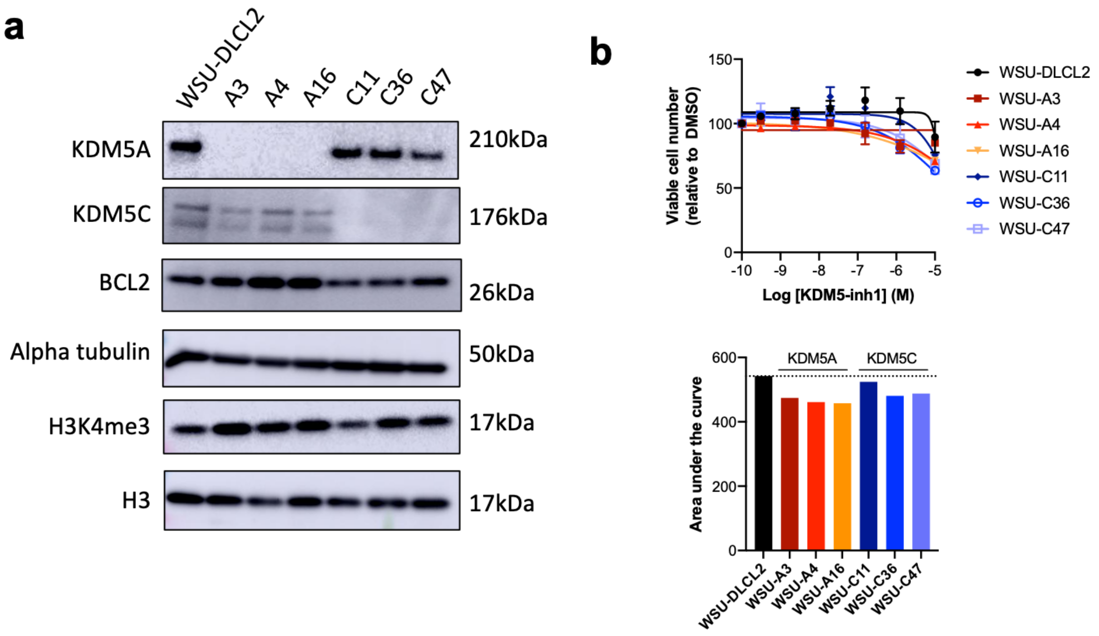


**Supplementary Figure 13. Knockout of *KDM5A* or *KDM5C* sensitizes cells to KDM5-inhibition. (a)** Western blot showing loss of KDM5A and KDM5C in three *KDM5A* homozygous knockout clones (WSU-A3, A30, A50) and three *KDM5C* homozygous knockout clones (WSU-C11, C36, C47), alongside expression of BCL2 and H3K4me3 levels. **(b)** WSU-DLCL2 cells and *KDM5A/KDM5C* knockout clones were exposed to DMSO or increasing concentrations of KDM5-inh1 for five days, and viable cell numbers quantified. Concentration response curves are shown in the upper panel and AUC values in the lower panel. Data are the mean ± SEM of three independent experiments.

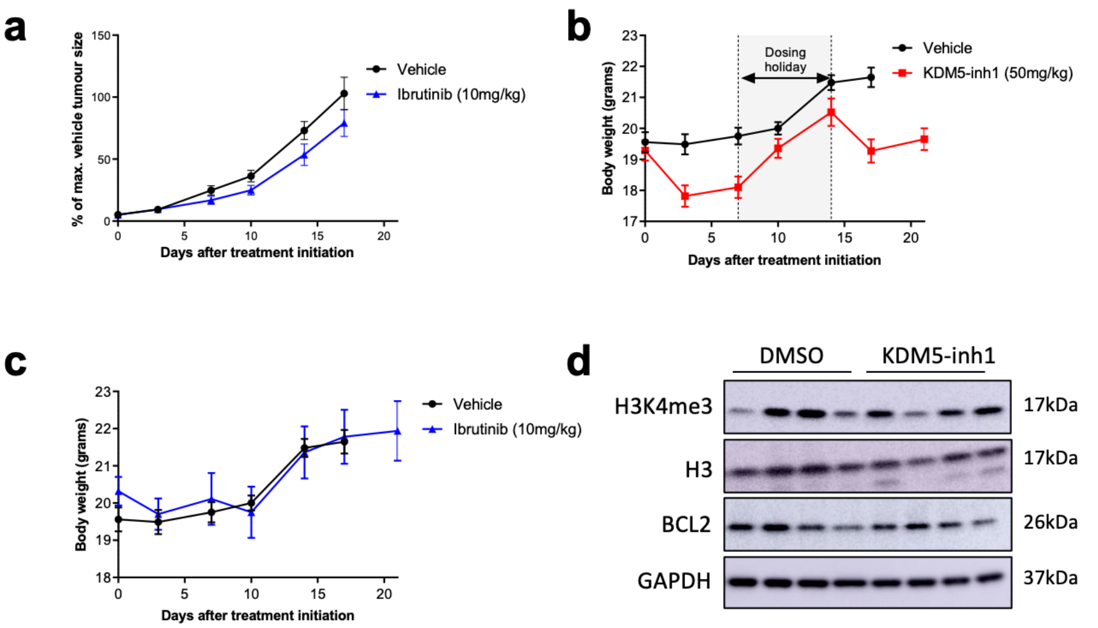


**Supplementary Figure 14.** ***In vivo*** **analysis of KDM5i. (a)** Activity of 10mg/kg ibrutinib on the growth of SU-DHL-6 xenografts, in comparison to vehicle treated mice. Effect of **(b)** 50mg/kg KDM5-inh1 and **(c)** 10mg/kg ibrutinib on the body weight of mice, in comparison to vehicle treated mice. Data are the mean ± SEM of 10 individual mice, except in the vehicle group where one mouse was removed due to insufficient tumour growth (<300mm$^3$). **(d)** Levels of H3K4me3, H3, BCL2 and GAPDH were quantified by western blot in tumours from mice treated with vehicle or 50mg/kg KDM5-inh1 for three weeks (including dosing holiday).

#### Supplementary Tables:

**Supplementary Table 1. KMT2D, KDM5A and KDM5C clones generated by CRISPR**

| Gene | Parental cell | Clone ID | Allele 1 | Allele 2 | TA cloned? |
|---|---|---|---|---|---|
| KMT2D | WSU-DCLCL2 | #8 | c.245_284del (40bp); L82Qfs*35 | WT | Yes |
| KMT2D | WSU-DCLCL2 | #22 | c.284_284delC; P95Qfs*35 | WT | Yes |
| KMT2D | WSU-DCLCL2 | #61 | c.245_284del (40bp); L82Qfs*35 | WT | Yes |
| KMT2D | HT | #4 | c.245_283del (39bp); L82_C94del | WT | No |
| KMT2D | HT | #9 | c.245_284del (40bp); L82Qfs*35 | WT | No |
| KMT2D | HT | #14 | c.244_257del (144bp); L82Vfs*4 | c.245_284del (40bp); L82Qfs*35 | No |
| KMT2D | HT | #16 | c.243-244insG; L82fs*5 | WT | No |
| KMT2D | HT | #18 | c.245_272del (28bp); L82Rfs*39 | c.245_284del (40bp); L82Qfs*35 | No |
| KMT2D | HT | #21 | c.244_305del (62bp); L82Pfs*5 | WT | No |
| KMT2D | SU-DHL-8 | K51 | c.1940_1941insC; P648fs*2 | WT | Yes |
| KMT2D | SU-DHL-8 | K65 | c.1940_1941insC; P648fs*2 | WT | Yes |
| KMT2D | SU-DHL-8 | K67 | c.1940_1941insC; P648fs*2 | WT | Yes |
| KDM5A | SU-DHL-6 | A3 | c.1404_1404delG; P469Rfs*49 | c.1404_1404delG; P469Rfs*49 | No |
| KDM5A | SU-DHL-6 | A30 | c.1404_1404delG; P469Rfs*49 | c.1404_1404delG; P469Rfs*49 | No |
| KDM5A | SU-DHL-6 | A50 | c.1404_1404delG; P469Rfs*49 | c.1404_1404delG; P469Rfs*49 | No |
| KDM5A | WSU-DLCL2 | A3 | c.1753_1769del(17bp); S464Vfs*16 | c.1753_1769del(17bp); S464Vfs*16 | No |
| KDM5A | WSU-DLCL2 | A4 | c.1767_1767delG; P469Rfs*49 | c.1767_1767delG; P469Rfs*49 | No |
| KDM5A | WSU-DLCL2 | A16 | c.1768_1772del (5bp); P469Afs*15) | c.1768_1772del (5bp); P469Afs*15) | No |
| KDM5C | WSU-DLCL2 | C11 | c.1674_1675del (2bp); E381Ifs*9 | c.1674_1675del (2bp); E381Ifs*9 | No |
| KDM5C | WSU-DLCL2 | C36 | c.1669_1678del (1bp); S379Lfs*14 | c.1669_1678del (1bp); S379Lfs*14 | No |
| KDM5C | WSU-DLCL2 | C47 | c.1141_1200+68del (128bp); E381_E400del | c.1141_1200+68del (128bp); E381_E400del | No |

**Supplementary Table 2. Supplementary Table 2. H3K4me3/H3K4me1 ChIP-seq analysis**

**Supplementary Table 3. RNA-seq analysis**

**Supplementary Table 4. GSEA analysis KDM5-inh1 treated cell lines**

**Supplementary Table 5. H3K4me1, H3K4me3 and RNA pathway analysis in WSU-DLCL2 and WSU#22 cells**

**Supplementary Table 6. Pathway analysis of SUDHL6 ChIP-seq data**

**Supplementary Table 7. Summary of drug combination**

| DrugA | DrugB | Cell_line | CSS | S | ZIP | BLISS | LOEWE | HSA |
|---|---|---|---|---|---|---|---|---|
| BCL2i | KDM5i | WSU-DLCL2 | 45.75 | -118.19 | -17.51 | -38.87 | -9.11 | -31.74 |
| BCL2i | KDM5i | KARPAS-422 | 53.2 | -95.58 | -20.23 | -43.42 | -23.35 | -30.38 |
| BCL2i | KDM5i | SUDHL6 | 54.31 | -109.59 | -17.05 | -35.02 | -14.09 | -31.43 |
| Ibrutinib | KDM5i | WSU-DLCL2 | 31.61 | -159.72 | -14.33 | -51.56 | -43.05 | -62.94 |
| Ibrutinib | KDM5i | KARPAS-422 | 38.62 | -75.59 | -24.73 | -55.45 | -25.81 | -39.63 |
| Ibrutinib | KDM5i | SUDHL6 | 28.42 | -106.4 | -22.67 | -52.79 | -11.61 | -48.83 |
| MCL1i | KDM5i | WSU-DLCL2 | 33.75 | -111.16 | -21.49 | -49.85 | -14.56 | -40.3 |
| MCL1i | KDM5i | KARPAS-422 | 21.49 | -75.66 | -29.28 | -62.77 | -45.88 | -55.14 |
| MCL1i | KDM5i | SUDHL6 | 28.38 | -104.44 | -27.96 | -60.16 | -12.61 | -55.94 |

**Supplementary Table 8. Details of cell lines used in study.**

| Cell line | Source | Disease | Growth medium | Mutation profiling | Cosmic exome available? | KMT2D allele 1 (amino acid) | KMT2D allele 2 (amino acid) | t(14;18) |
|---|---|---|---|---|---|---|---|---|
| OCI-LY-1 | DSMZ | GCB-DLBCL | 20% FBS, IMDM | Targetted | No | P4929Lsf*66 | R1903* | Yes |
| OCI-LY-7 | DSMZ | GCB-DLBCL | 20% FBS, IMDM | Targetted | Yes | WT | WT | No |
| OCI-LY-18 | DSMZ | GCB-DLBCL | 10% FBS, RPMI | Exome | No | P658fs | splice site disrupted (chr12, 49446988; A>C) | Yes |
| SU-DHL-4 | Tissue bank | GCB-DLBCL | 10% FBS, RPMI | Targetted | Yes | WT | WT | Yes |
| SU-DHL-6 | Tissue bank | GCB-DLBCL | 10% FBS, RPMI | Targetted | Yes | E4712* | Q211* | Yes |
| SU-DHL-8 | DSMZ | GCB-DLBCL | 10% FBS, RPMI | NA | Yes | P648Tfs*2 | P648fs*2 | Yes |
| KARPAS-422 | Tissue bank | GCB-DLBCL | 10% FBS, RPMI | Targetted | Yes | Q3278* | WT | Yes |
| DOHH2 | Tissue bank | GCB-DLBCL | 10% FBS, RPMI | Targetted | Yes | WT | WT | Yes |
| HT | DSMZ | GCB-DLBCL | 10% FBS, RPMI | Targetted | No | WT | WT | No |
| WSU-DLCL2 | DSMZ | GCB-DLBCL | 10% FBS, RPMI | Targetted | Yes | WT | WT | Yes |
| OMP2 | Tissue bank | Multiple myeloma | 10% FBS, RPMI | NA | Yes | G1486D | Q3322* | No |
| RAJI | Tissue bank | Burkitt's lyphoma | 10% FBS, RPMI | NA | Yes | WT | WT | No |
| WSU-NHL | DSMZ | GCB-DLBCL | 10% FBS, RPMI | NA | Yes | V2281fs*32 | L2425fs*26 | Yes |
| DB | DSMZ | GCB-DLBCL | 10% FBS, RPMI | NA | Yes | Q2736* | P480fs* | Yes |

**Supplementary Table 9. Antibodies used in study.**

| Target | Supplier | Product no. |
|---|---|---|
| phospho-SYK (Tyr525/526) | Cell Signaling Technologies | 2710 |
| SYK | Cell Signaling Technologies | 2712 |
| GAPDH | Cell Signaling Technologies | 2118 |
| alpha-Tubulin | Sigma Aldrich | T9026 |
| H3K4me3 (for western blots) | Cell Signaling Technologies | 9751 |
| H3K4me2 (for western blots) | Cell Signaling Technologies | 9725 |
| H3K4me1 (for western blots) | Cell Signaling Technologies | 5326 |
| H3 | Cell Signaling Technologies | 4499 |
| H3K9me3 | Active Motif | 39161 |
| H3K36me3 | Diagenode | C15200183 |
| H3K27me3 | Cell Signaling Technologies | 9733 |
| H3K27ac | Active Motif | 39133 |
| H3K4me3 (for ChIP) | Diagenode | C15410003 |
| H3K4me2 (for ChIP) | Diagenode | C15200151 |
| H3K4me1 (for ChIP) | Diagenode | C15410194 |
| Spike-in Antibody | Active Motif | 61686 |
| KDM5A | Abcam | ab70892 |
| KDM5B | Sigma Aldrich | HPA027179 |
| KDM5C | Abcam | ab34718 |
| KDM5C | Diagenode | C15410338 |
| KDM5D | Sigma Aldrich | HPA049086 |
| PTPN6 | Santa Cruz | sc-7289 |
| BCL2 | BD sciences | 610538 |
| PARP | BD sciences | 556494 |
| MCL-1 | Santa Cruz | sc-819 |
| BCL-XL | Cell Signaling Technologies | 2764 |
| BCL-2 | DAKO | M0887 |
| NOXA | Merck millipore | OP180 |
| BIM | Cell Signaling Technologies | 2933 |
| HSC70 | Santa Cruz | sc-7298 |
| Goat F(ab’)2 Anti-Human IgG-UNLB | Cambridge Bioscience | 0110-01 |
| Goat F(ab’)2 Anti-Human IgM-UNLB | Cambridge Bioscience | 2022-01 |
| Anti-mouse | DAKO | P0448 |
| Anti-Rabbit | DAKO | P0447 |

**Supplementary Table 10. Primers used in study.**

**qRT-PCR**

| Target | Forward | Reverse |
|---|---|---|
| KMT2D | CCCCTGAGAGCTGCTGTG | GTAACGGGTGATGGGCAAAA |
| BCL2 | GGTGGGGTCATGTGTGTGG | CGGTTCAGGTACTCAGTCATCC |
| BCL2A1 | TACAGGCTGGCTCAGGACTAT | CGCAACATTTTGTAGCACTCTG |
| MYB | GAAAGCGTCACTTGGGGAAA A | TGTTCGATTCGGGAGATAATTG G |
| CD72 | ATCTGAGGTTTGTGAAGGCTC C | AACATTCTCGTAGGTGATTTCCC |
| FCGR2B | AGCCAATCCCACTAATCCTGA | GGTGCATGAGAAGTGAATAGGT G |
| FCRL5 | ACCCAGGCCCATTATTTTCCT | AGTAGAAGCGAAATCCCTTGC |
| PTPN6 | GGAGAAGTTTGCGACTCTGA C | GCGGGTACTTGAGGTGGATG |
| DUSP6 | GAAATGGCGATCAGCAAGAC G | CGACGACTCGTATAGCTCCTG |
| PLCB2 | ATCCGGGATACTCGCTTTGG | CACCACCGTGAGTGTCTTCAG |
| KDM5A | AGCCGAGTTGGGAGGAGTT | TGGACTCTTGGAGTGAAACGA |

**ChIP-PCR**

| Target | Forward | Reverse |
|---|---|---|
| PTPN6 (537bp) | TTCTCGCTCTCCGTCAGGTA | GGGGAACCAGGAATGAGTGG |
| PTPN6 (827bp) | ATGGAGGGGAGAAGTTTGCG | GGAGCCCTCACCTCTCACTA |
| PTPN6 (-634bp) | GCTCAGGGTCATGTTGTCCA | GGTGCCCTCCTCTAACCAAC |
| PTPN6 (3016bp) | CATCCGCCTTCCTTGTGACT | AAGAGAAACGCAGACCGAGG |
| FCRL5 promoter | GCCCATGGTGAGCCCTTTTA | CCTTGGTGGTCCAGGTCTTC |
| DUSP6 promoter | ATTCGGACTCCGTGCTACTG | AAGACGCCCGGGTAGATTTG |
| FCGR2B promoter | AGAGAGGAACGGGAACCTCA | ATCTTCACCAGCCTGCCTTC |
| PLCB2 promoter | AAAGGTTGGGAGCACCGTAG | AAGAGGAGCCGTGTGTTCAG |
| CD72 promoter | GCTCTCCAGACCTGCTTTGT | GAATGTTCAAGTGCCCGCAG |
| BCL2 promoter (-6bp) | AATGAATCAGGAGTCGCGGG | GGGATTCCTGCGGATTGACA |
| BCL2 promoter (-1558bp) | ATCCACAGGGCGATGTTGTC | GAACTGGGGGAGGATTGTGG |
| BCL2 promoter (523bp) | CTGCTACGAAGTTCTCCCCC | ACCAGGAGGAGGAGAAAGGG |

**Supplementary Table 11. guideRNAs used in study.**

| ID | Sequence |
|---|---|
| KMT2D gRNA 1 | TGATTGGCCCCGGTGTCCAG |
| KMT2D gRNA 2 | AGTTGCCATTTGATTGGCCC |
| KMT2D gRNA 3 | TACACGGGCAGCGGGAGCTA |
| KMT2D gRNA 4 | CGTTGTGCTCTCTGTAACTG |
| SU-DHL-8 KMT2D gRNA 1 | TAGGCGCGATACCTCAGGTG |
| SU-DHL-8 donor template | TCCCCACCTGAGGACTCGCCTATGTCCCCACCACC<br>TGAAGAATCACCTATGTCCCCACCACCTGAGGTAT<br>CGCGCCTATCCCCCCTGCCTGTGGTGTCACGCCTGT<br>CTCCACCGCC |
| KDM5A gRNA 1 | TCTCTGGTATGAAAGTGCCG |
| KDM5C gRNA 1 | GAAACCGCTGCCAAATTCTT |

**Supplementary Table 12. Transfection conditions.**

| Cell Line | Instrument | Buffer | Program | Cell number |
|---|---|---|---|---|
| WSU-DLCL2 | Nucleofector II | Buffer T | G-016 | 5 million |
| HT | Nucleofector II | Buffer T | A-023 | 5 million |
| SU-DHL-8 | Nucleofector II | Buffer T | A-032 | 5 million |
| SU-DHL-6 | Nucleofector 4D | Buffer SF | CA-137 | 1 million |
